# 3-dimensional arenas for the assessment of *C. elegans* behavior

**DOI:** 10.1101/2021.11.11.468110

**Authors:** Steel N. Cardoza, Lai Yu Leo Tse, Kira Barton, Eleni Gourgou

## Abstract

*C. elegans* nematodes are a well-established model organism in numerous fields of experimental biology. In nature, *C. elegans* live in a rich 3-dimensional environment. However, their behavior has been assessed almost exclusively on the open, flat surface of NGM (Nematode Growth Medium) plates, the golden standard for *C. elegans* culture in the lab. We present two methods to build 3-dimensional behavioral arenas for *C. elegans*, by casting, and by directly 3D printing NGM hydrogel. The latter is achieved by using a highly customized fused deposition modeling (FDM) 3D-printer, modified to employ NGM hydrogel as ink. The result is the advancement of 3-dimensional complexity of behavioral assays. To demonstrate the potential of our method, we use the 3D-printed arenas to assess *C. elegans* physical barriers crossing. *C. elegans* decision to cross physical obstacles is affected by aging, physiological status (i.e., starvation), and prior experience. The 3D-printed structures can be used to spatially confine *C. elegans* behaviors, i.e., egg laying. We consider these findings a decisive step toward characterizing *C. elegans* 3-dimensional behavior, an area long overlooked due to technical constrains. We envision our method of 3D-printing NGM arenas as a powerful tool in behavioral neurogenetics, neuroethology, and invertebrate model organisms’ neurobiology.

## Introduction

*C. elegans* nematodes are a well-established model organism in numerous fields of experimental biology, with prominent among them the biology of aging, behavioral neurogenetics, and neurobiology (1–4). In nature, *C. elegans* live in a rich 3-dimensional environment (e.g., rotten fruit, muddy soil) (5). However, their behavior has been assessed almost exclusively on the open, flat surface of NGM (Nematode Growth Medium) plates (6), which are the golden standard of *C. elegans* culture in the lab.

Recently, a new type of associative learning was reported (7), observed in T-shaped mazes (Fig. 1A). *C. elegans* learning to reach a target T-maze arm is related to the 3-dimensional nature of the arena (7, 8), namely walls, floor, and overall surfaces, which are perceived through multiple sensory modalities. In addition, *C. elegans* show a clear preference for richly patterned surfaces (9). Combined, these findings support the idea that 3-dimensional environments are required to witness the full behavioral expression of these nematodes, and a larger part of their nervous system’s capacity. *C. elegans* 3-dimensional locomotion is also gaining a lot of attention (10–13), revealing very interesting dynamics in both swimming and crawling nematodes.

**Figure 1:**
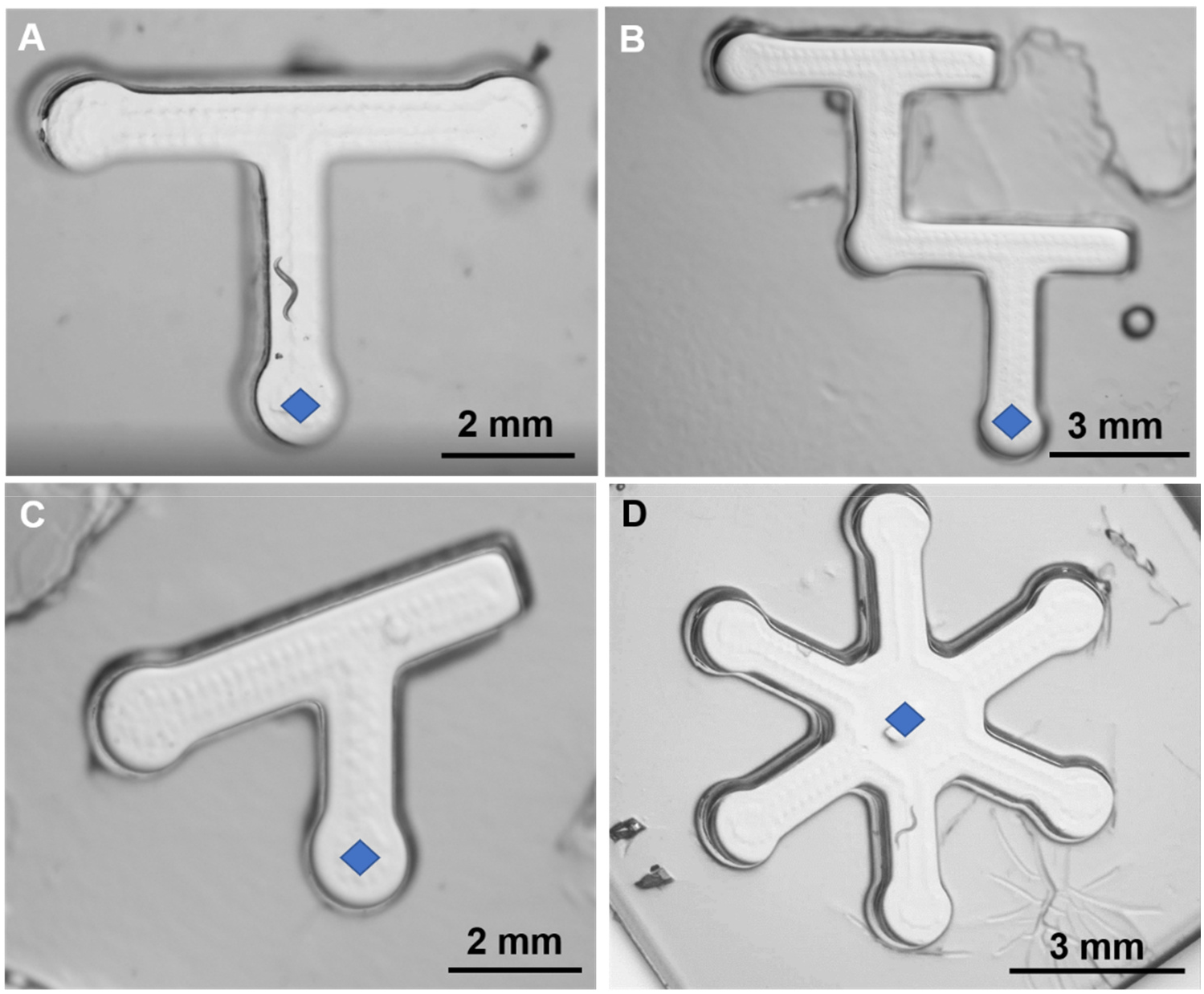
NGM behavioral arenas for *C. elegans,* made with molding technique. (**A**) A simple, symmetrical T-maze (see also Gourgou et al, 2020). (**B**) Double T-maze, a more complex version of T-maze. (**C**) Angled T-maze with asymmetrical ends. (**D**) Radial arena, with six arms. All panels: mazes/arenas depth: ∼5 mm; diamond shape indicates the intended assay starting point.

To explore *C. elegans* spatial behavior, our group uses customized arenas (Fig. 1). To this end, 3D-printed plastic molds are used to imprint mazes of various designs in NGM undergoing solidification (7, 8). This leads to reliable, highly repeatable, and importantly, nematode-friendly assays because the generated arenas are built entirely out of NGM (Fig 1). However, several constrains apply. Although the complexity in the *x,y* dimensions (plane parallel to the assay plate surface) can increase freely (Fig. 1A-D), complexity in *z* dimension (plane perpendicular to the assay plate surface) is essentially limited to varying the depth of the arenas. Vertical elements, suspended features, multi-level structures and other similar architectures are not allowed using molds.

To meet the need for alternative fabrication processes, we explored two methods. The first one includes the use of PVA (polyvinyl alcohol), a water-soluble synthetic polymer, to cast NGM structures (Figs 2 and 3). The ensuing parts are of high quality and provide valuable feedback regarding the NGM self-sustaining properties. However, this method’s limitations motivated us to seek another route, one that employs a 3D printer, which uses NGM as ink.

**Figure 2:**
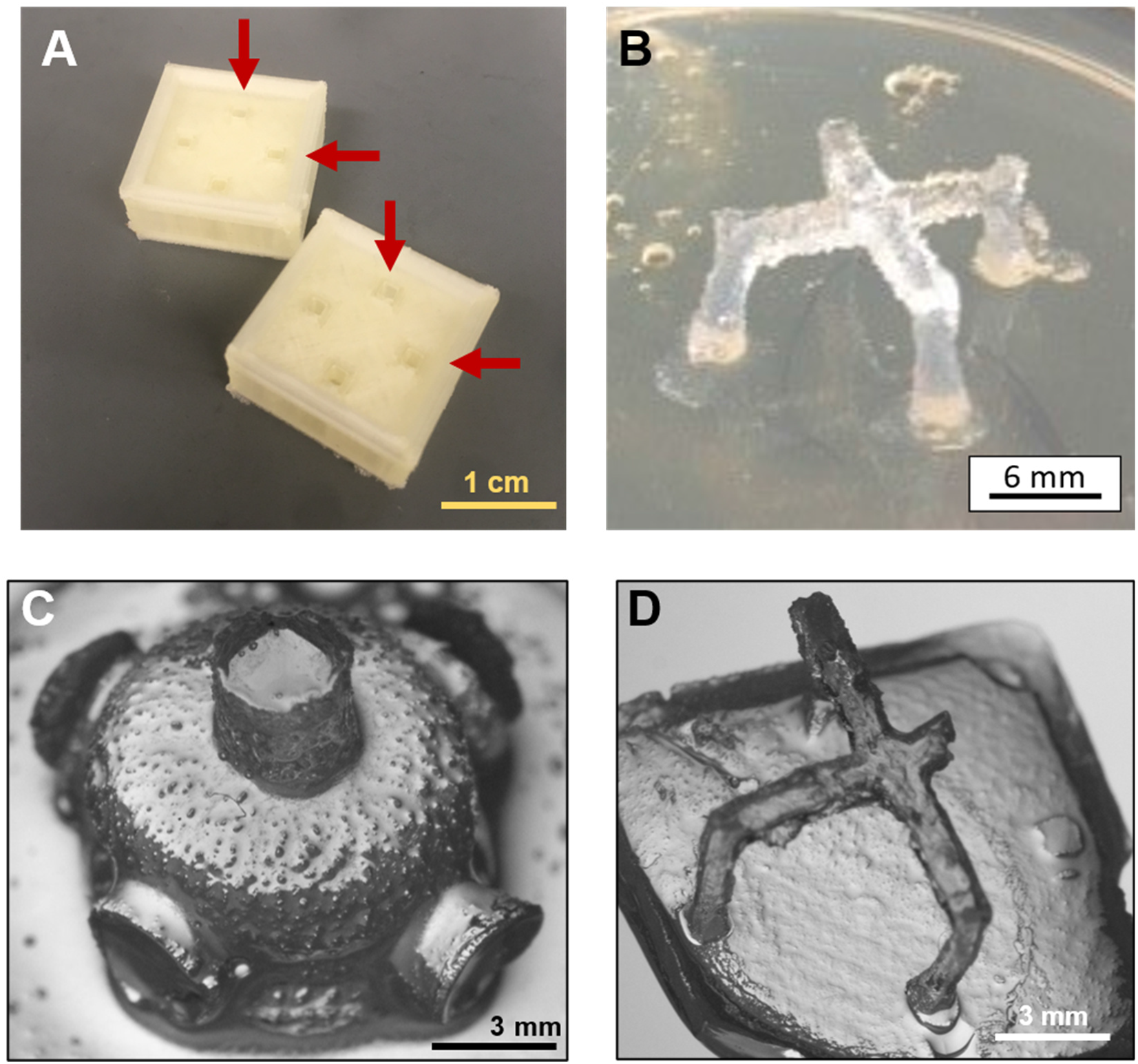
NGM 3-dimensional structures, made with the PVA casting method. (**A**) Two 3D-printed PVA casting molds, red arrows indicate the liquid NGM input point. (**B**) A four-legged NGM crossbridge, made using one of the casts shown in (A), sitting on a NGM plate surface. Legs are slightly tilted outwards because of flipping and handling the structure. (**C**) A diving bell – like structure, consisting of a hemisphere (radius: 5mm) and designed to have five cylindrical arms, diameter: 3mm, length: 1-1.5mm, each. Liquid NGM did not reach the entire length of the hollow space inside the PVA cast, resulting in much shorter arms than originally planned (4mm). Note the rough surface of the structure. (**D**) A four-legged crossbridge standing on a 2mm thick base, raised 4 mm above the base’s surface. This is a much thinner structure than the one in (B), showcasing the self-supporting properties of NGM even in smaller arrangements. Note the missing right arm.

**Figure 3:**
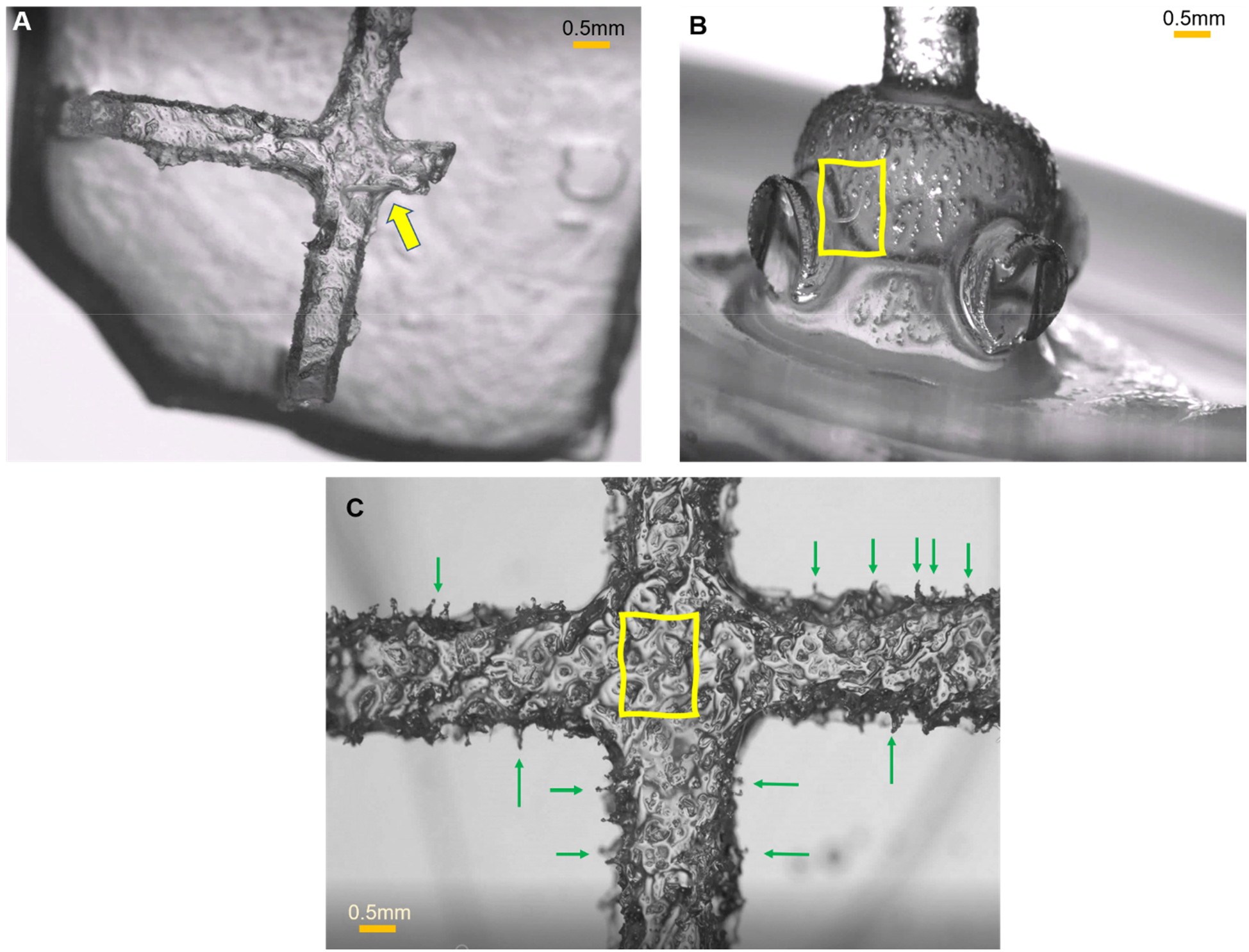
Properties of PVA-casted 3-dimensional NGM structures. (**A**) Closeup of the four-legged NGM crossbridge, shown in Fig. 2D, top view. Yellow arrow points to a *C. elegans* nematode crawling on the upper surface, at the corner between two beams. Note the rough surface and the thin, relatively to worm’s body length, structure. (**B**) Closeup of the NGM diving bell-like structure, shown in Fig. 2C, side view. Yellow frame indicates the position of a *C. elegans* nematode. Note the bumpy surface and the incomplete beams. (**C**) Closeup of the four-legged NGM crossbridge, shown in Fig. 2B, top view. Yellow frame indicates the position of a *C. elegans* nematode. Note the very rough surface, of which the protruding features (examples highlighted with green arrows) are similar to or even bigger than the worm’s body width. The *C. elegans* worm is challenging to distinguish in all three photos.

A growing number of researchers are employing 3Dprinting as a transformative tool for cell and tissue engineering (14, 15). This includes 3-dimensional scaffolds made of enriched hydrogel-based materials (16). Hydrogels of 1% −5% agar concentrations have been successfully explored for 3D bioprinting applications (17). Most of the occurring structures are sturdily self-supported cubes or other non-hollow, no-overhang designs. Interestingly, the 3D-printing technology has not been used to produce behavioral arenas for the study of small invertebrate animal models, like *C. elegans*.

We present a highly customized prototype 3D-printer, the Parnon Printer (Parnon: mountain in southern Greece, known for its many gorges), which can print 3-dimensional parts, suitable for *C. elegans* behavioral experiments, made of 2% agar-based NGM hydrogel. The resulting arenas are nematode-friendly, minimizing the stress that could have been induced when the animals are transferred from the culture plate into the arenas.

To demonstrate the suitability of the Parnon-printed parts, we use them to assess *C. elegans* physical barrier crossing ability, in the context of aging (young, middle-aged adults), feeding history (fully fed, starved animals) and prior experience (have been or not in the presence of a 3D structure before). We also explore the usage of 3D-printed structures to spatially confine *C. elegans* egg laying behavior. *C. elegans* behavior in 3-dimensional environments is by definition not possible to explore on standard flat NGM plates. Therefore, the findings reported here would likely not have been brought to light if the Parnon Printer had not been developed.

## Results

### PVA casted 3D-structures

The first fabrication method explored was PVA casting (Fig. 2). The produced structures show that it is possible to create 3-dimensional NGM structures in a systematic and predictable way.

Specifically, the crossbridge design showed that NGM can support itself, even at a height of 4 mm and an overhang length of 12 mm, with legs 2×2mm in cross section(Fig 2B). Self-standing crossbridges are also feasible in smaller structures (Fig. 2D), 2mm high and 5mm long overhang, where the bridge legs are thinner, 1.5mm in cross section. This means that the mechanical properties of NGM 2% in agar are such that allow for overhangs which stretch a few worm body lengths long, without the need of extra support. It is noted that during the solidification process, NGM enjoyed the support provided by the PVA cast itself (Fig. 2A).

Cross sectional dimensions of beams to as small as 1×1mm have been attempted but were found too small to consistently allow NGM to enter all the way into the cast channels (Fig 2D, missing right arm). This was similarly the case when casting the diving bell design (Fig. 2C, 3B), where the cross section of the arms was also 1×1mm.

PVA casting produces parts with very rough surfaces (Fig. 3). This is a consequence of the cast 3D-printing process, during which the PVA is laid in a way that allows micropockets of air among the deposed PVA threads. These micropockets get filled later with NGM, thus creating tiny protrusions (Fig. 3C, green arrows).

### Parnon customization and Parnon-printed 3D-structures

We extensively modified the commercially available FDM (Fusion Deposition Modeling) printer of choice (Fig. 4, Supplementary Fig. S1) and converted it in a highly customized hydrogel-ink 3D printer, named Parnon. Customization included wide-ranging modifications of the print head and substrate (Fig. 4C, 4D, Supplementary Figs. S2, S3A), and involved design, 3D-printing (motor arm housing, syringe plunger connector), and machining (print head aluminum heat sink) of tailored parts (Supplementary Fig. S1). Effective synergy of all parts was imperative for the successful operation of the instrument.

**Figure 4:**
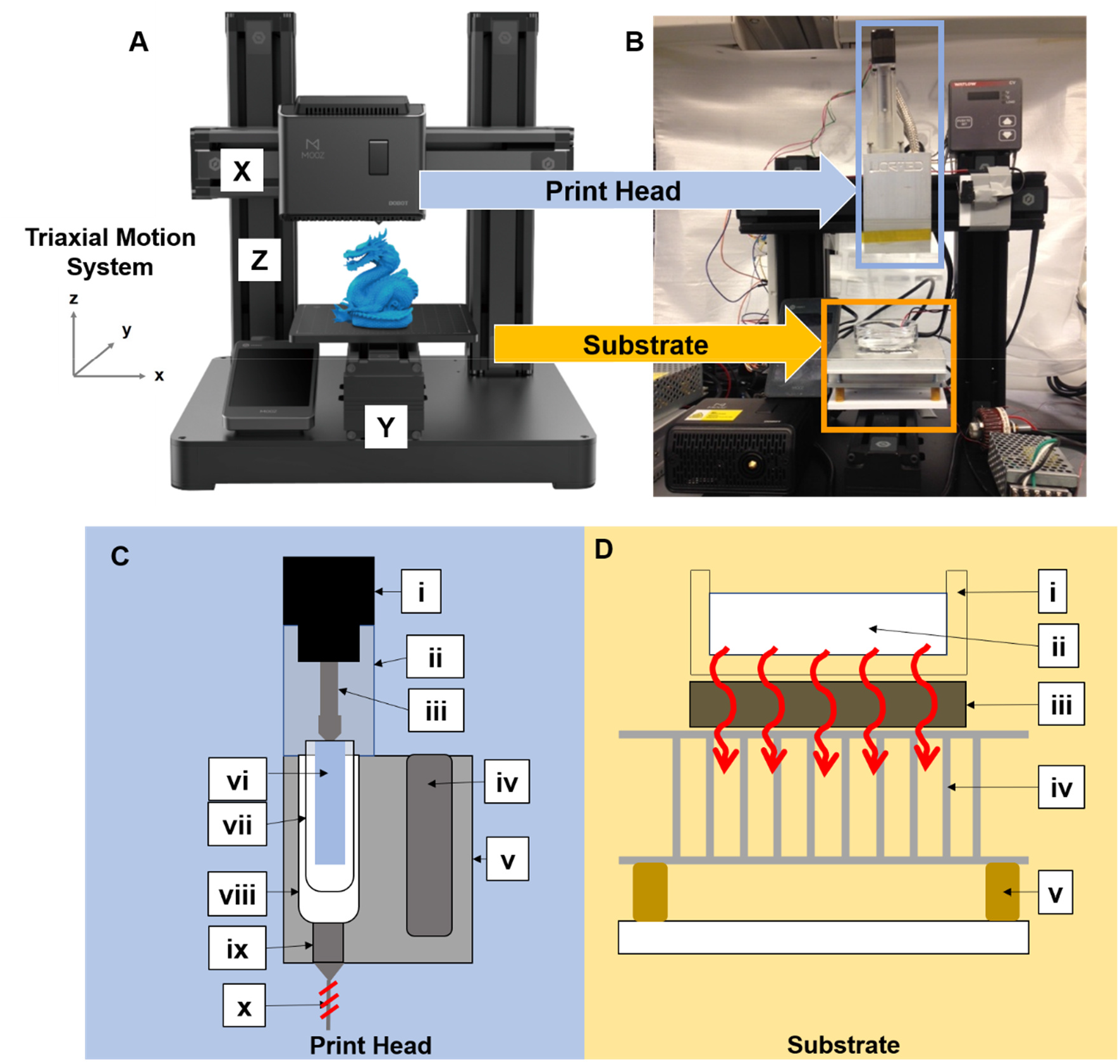
Overview of customization changes that convert a FDM printer to Parnon Printer. (**A**) The FDM 3D Printer DOBOT MOOZ (image courtesy of the manufacturer). The three main parts are the print head, the substrate, and the tri-axial relative motion system. While the stock motion system was considered adequate, both the print head and substrate had to be modified. (**B**) The prototype Parnon Printer, featuring a customized print head and substrate, designed for printing NGM. See also Supplementary Fig. S1. (**C**) Customized print head, front view schematic, connected to the x axis of the triaxial motion system. i) Bipolar stepper motor of the linear actuator. ii) 3D-printed custom connector, houses the stroke arm (iii) and connects the motor (i) to the heat sink (v). iii) Stroke arm of the linear actuator. iv) Heating element, heats the aluminum sink, keeping NGM in liquid state. v) Custom aluminum heat sink. vi) Custom connector connects the stroke arm (iii) to the plunger (vii). vii) Glass syringe plunger adhered on the custom connector. viii) Glass syringe houses the liquid NGM. ix) Metal luer-lock nozzle. x) Copper wire heat induction system, to prevent NGM solidification in the nozzle (ix). Details on the print head parts in Appendix. (**D**) Customized substrate, front view schematic, connected to the Y axis of the triaxial motion system. i) Glass petri dish. ii) Plotting medium. iii) Peltier device, red arrows indicate direction of heat transfer. iv) Heat sink to promote the thermal gradient created by the Peltier device (iii). iv) Springs used as a four-point bed leveling system (Supplementary Fig. S2).

Parnon printer can successfully use NGM as ink to print 3-dimensional structures (Fig. 5 and Suppl. Fig. 7). Although the examples presented here are of lower complexity compared to PLA and other plastic or resin 3D-printed objects, essential properties of 3D-printed structures are achieved. Hence, the Parnon-printed parts consist of multiple layers (up to 3), making this is an effective way to increase vertical complexity of behavioral arenas, in the dimension perpendicular to the surface of NGM plates. More layers are mechanically feasible, however currently Parnon does not allow for very precise stacking of deposed layers, which results in the top layers and bottom layers being misaligned (Fig. 5B).

**Figure 5:**
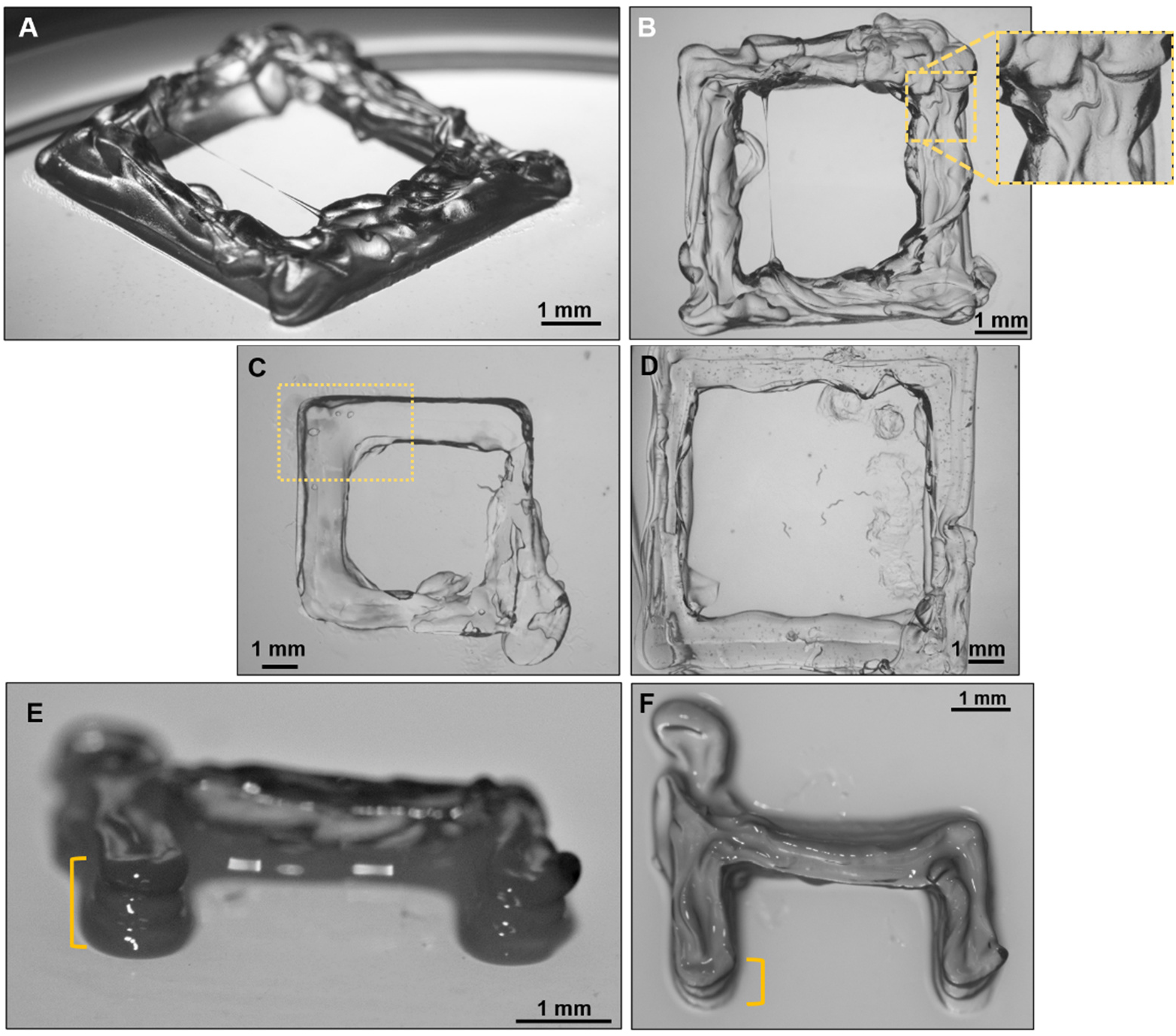
Parnon-printed NGM 3-dimensional structures. The Parnon-printed 3D structures have visibly smoother surface compared to cast-made ones (see Fig. 3). (**A**) Perspective and (**B**) top view of a 7×7mm square-shaped print, consisting of 3 layers, 500*μm* thick each. Printing conditions: head speed 10mm/s, actuation speed 31*μm/s*, printing head specs: nozzle ID 813 *μm*, linear actuator NEMA 14. Inset: a clearly distinguishable nematode, crawling on the printed NGM. (**C**) Top view of a 7×7mm square print, consisting of 3 layers, 300 *μm* thick each. Printing conditions: head speed 10 mm/s, actuation speed 38*μm/s*, printing head specs: nozzle ID 254*μm*, linear actuator NEMA 8. Yellow frame indicates the smoothly deposed NGM line at the corner of the square (90° angle deposition). (**D**) Top view of a 20×20mm square print, consisting of 3 layers, 300 *μm* thick each. Printing conditions: head speed 10 mm/s, actuation speed 8*μm/s*, printing head specs: nozzle ID 254*μm*, linear actuator NEMA 8. (**E**) Perspective and (**F**) top view of a C-shaped print, consisting of 3 layers, 300 *0* thick each. Printing conditions: head speed 10 mm/s, actuation speed 38*μm/s*, printing head specs: nozzle ID 254*μm*, linear actuator NEMA 8. Although C-shaped parts were not used in any experiments, they are included here to demonstrate the layering details. Yellow bracket indicates the beveled alignment of layers. Parts in C, D, E, and F are printed with a higher resolution print head (combination of nozzle diameter and linear actuator properties), compared to the part in A & B. Prints from both print heads are shown for comparison. For reference, the squared-shaped prints shown in Figures 6 – 8 were generated using the higher resolution print head, as well (see specifications in Supplementary Information section). See also Supplementary Fig. 7.

Importantly, the Parnon-printed parts have smooth surfaces, especially when they are printed using a narrower nozzle and higher resolution printing head (Fig. 5A, 5B, 5E, 5F). The extrusion is unhindered and continuous during a 90° angle turn (example: Fig. 5E, framed area) and NGM lines can be laid successfully in 90° angles or smaller (Suppl. Fig. 7B).

### C. elegans ability to cross physical barriers, and the effect of aging

To confirm that Parnon-printed structures can be used to investigate *C. elegans* behavior, we ran a series of experiments. In the first scheme, we assessed nematodes’ ability to cross physical barriers, in order to reach a food source (Figs 6 and 7). We also examined the role of aging (Fig. 6), feeding history, and prior experience (Fig. 7).

**Figure 6:**
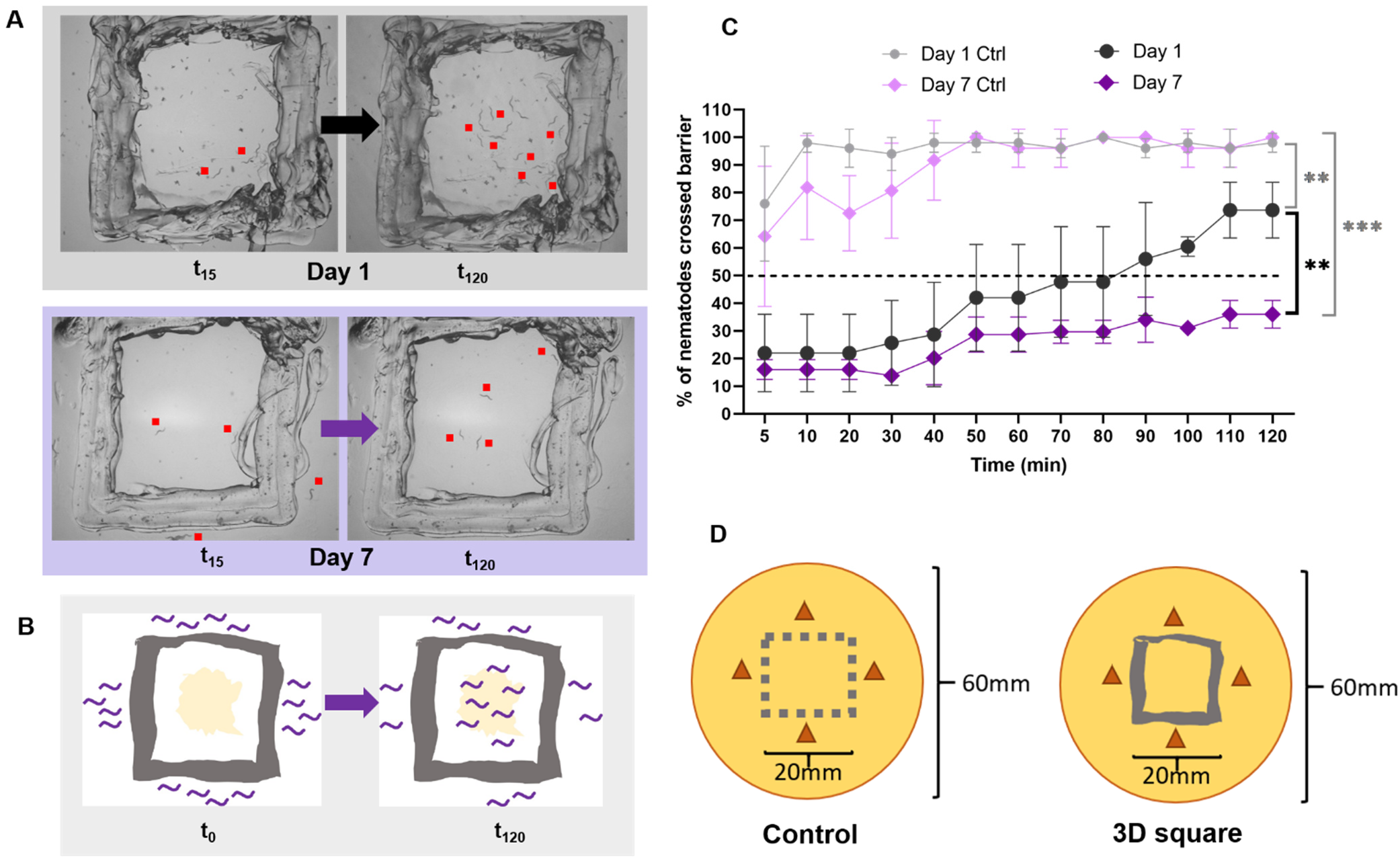
Assessment of *C. elegans* ability to cross a physical barrier to reach a food source. (**A**) Top panel: Snapshots of the square barrier assay when a population of adult Day 1 nematodes is tested, at t_0_ (left) and t_120_ (right); Bottom panel: Snapshots of the square barrier assay when a population of adult Day 7 nematodes is tested, at t_0_ (left) and t_120_ (right). Red dots indicate location of nematodes; number of dots does not correspond to number of nematodes. Scattered dark spots in the substrate are noticeable due to localized crystallization of NGM components, which often happens if the plates are a few weeks old (see Supplementary Information), however the overall suitability of the plate is not compromised. (**B**) Schematic of the assay. An NGM 3D-printed square (gray) is placed on an NGM plate, and *E. coli* OP50 food (yellow) is pipetted inside the square. A population of *C. elegans* nematodes (purple lines) is transferred on the NGM plate, outside of the square, at least 3mm away from it. At t_0_ no nematode is inside the square (nematodes placed on assay plate). At t_120_ a fraction of the worms’ population has crossed the square barrier and is foraging on the food. For the control experiment, the 3D-printed square is absent, and a figurative square is marked at the bottom of the plate, to define the target area (6D, left). (**C**) Graph showing the % of two age groups of nematodes that crossed the square barrier over 120min; purple diamonds: adult Day 1 hermaphrodites, (three independent experiments, n_1_=18, n_2_=12, n_3_=11); black circles: adult Day 7 hermaphrodites, (three independent experiments, n_1_=12, n_2_=16, n_3_=16). As a control experiment, adult day 1 and adult Day 7 nematodes were assayed on dishes with figurative square frame over 120min; pink diamonds: adult Day 1 hermaphrodites, (three independent experiments, n_1_=21, n_2_=17, n_3_=17); grey circles: adult Day 7 hermaphrodites, (three independent experiments, n_1_=6, n_2_=8, n_3_=9); Comparisons made using multiple unpaired t-tests, *p* values significant when *p*>0.05; **:*p*<0.01,***: p<0.001, only significant comparisons between final (at t=120min) are shown; error bars indicate standard deviation, dashed line indicates 50% level. See also Supplementary Table S1. (**D**) Schematic showing the relative size and position of the figurative square frame used in the control experiments (left) and of the 3D-printed square used in the physical barrier experiments (right), with respect to a 60mm petri dish. 3D-printed squares were ∼20×20mm, 3 layers, 0.5mm thick each. Brown triangles indicate initial placement of nematodes.

**Figure 7:**
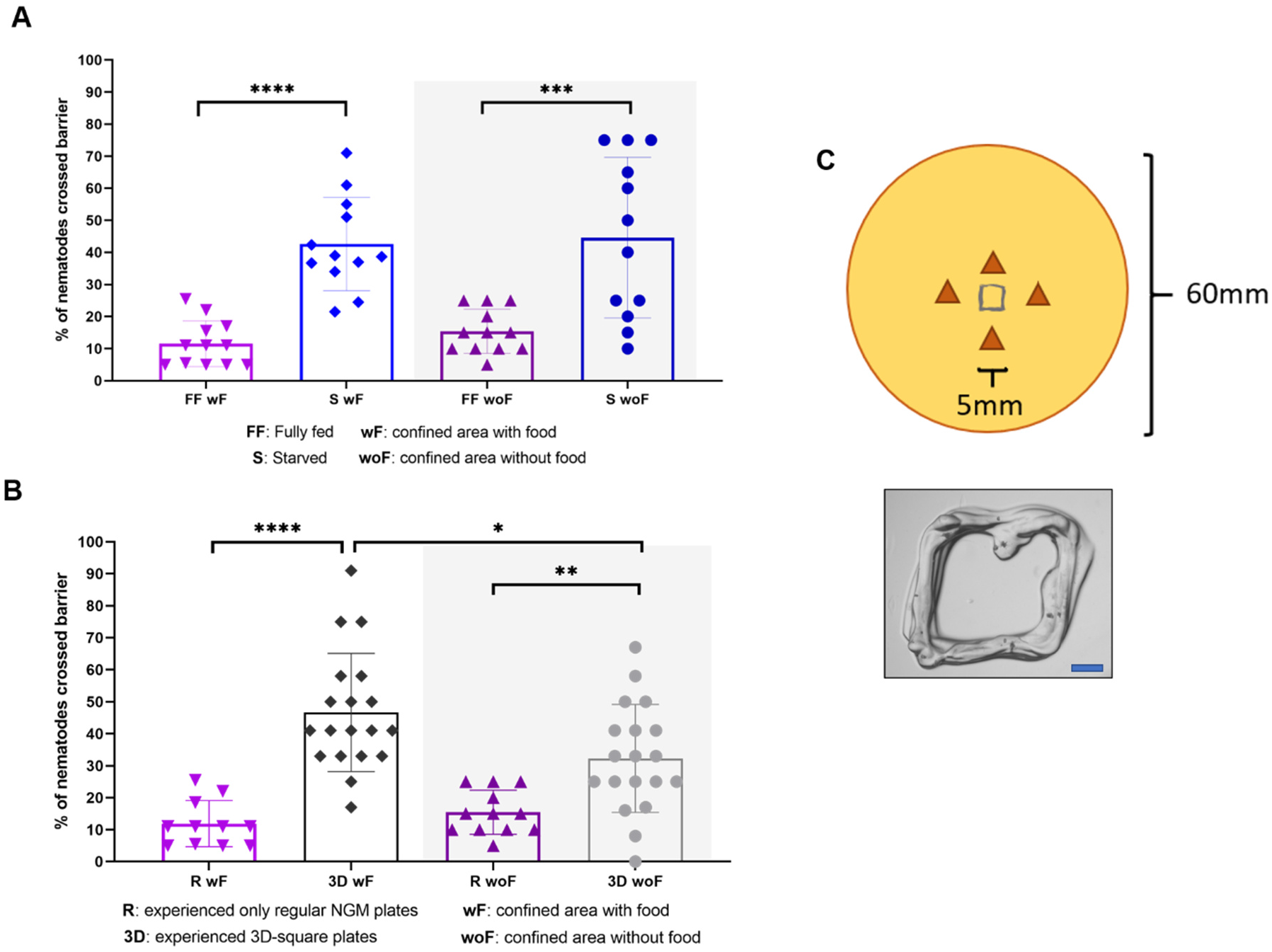
*C. elegans* physical barrier crossing with respect to age and feeding status. (**A**) Graph showing the % of four groups of adult Day 1 nematodes that crossed the square barrier over 120min; purple inverted triangles: nematodes that have been fully fed (FF), cross the barrier into a square baited with food (wF); blue diamonds: nematodes that have been starved (S) for 24 hours prior testing, cross the barrier into a square baited with food (wF); purple triangles: nematodes that have been fully fed (FF), cross the barrier into a square without food (woF); blue circles: nematodes that have been starved (S) for 24 hours prior testing, cross the barrier into a square without food (woF). (**B**) Graph showing the % of four groups of fully fed Day 1 adult nematodes that crossed the square barrier over 120min; purple inverted triangles: nematodes that have been grown on regular NGM plates (R), cross the barrier into a square baited with food (wF); black diamonds: nematodes that have experienced a 3D square for 24 hrs prior testing (3D), cross the barrier into a square baited with food (wF); purple triangles: nematodes that have been grown on regular NGM plates (R), cross the barrier into a square without food (woF); grey circles: nematodes that have experienced a 3D square for 24 hrs prior testing (3D), cross the barrier into a square without food (woF). Data of *C. elegans* grown on regular NGM plates (purple triangles and inverted triangles, R) are the same with fully fed nematodes data (FF) in panel A. Panels A and B: Each data point corresponds to the percentage of worms scored inside the target square at a given timepoint, measurements taken every 5 or 10min over a 120min period, horizontal lines indicate the mean, error bars indicate the standard deviation. Comparisons made using two-tailed, unpaired t-test, *p* values significant when *p*>0.05; *: *p*<0.05, **:*p*<0.01, ***: *p*<0.001, ****: *p*<0.0001, only significant comparisons shown. Shaded area: confined area did not contain food (without food: woF). (**C**) Top: Schematic showing the relative size of the 5×5mm 3D-printed square with respect to a 60mm petri dish, brown triangles indicate initial placement of nematodes. Bottom: Example of a 3D-printed square used in the above experiments, 5×5mm, 3 layers, 0.5mm thick each; scale bar: 1mm.

To establish that the 3D printed squares constitute a physical barrier for *C. elegans*, we conducted a control experiment, in which adult Day 1 nematodes are allowed to reach a food source not framed by a physical barrier (Fig. 6B, and 6D, left). Animals reached the food containing area in large numbers quickly, and almost all of them (96-100%) remained there for most of the 120min assay (Fig. 6C, grey circles).

Next, we challenged adult Day 1 *C. elegans* with a food source, framed by a 20×20mm 3D-printed NGM square (Fig. 6D, right). In this case, nematodes entered the framed area gradually and in lower rates, and at the end of the 120min assay, 74% of animals had crossed the square barrier and had reached the food source (Fig. 6C, black circles). Therefore, the NGM square frame presents a physical barrier for nematodes.

We hypothesized that decision making related to physical challenges differs in young and old animals, as aging-driven behavioral changes have been broadly reported in *C. elegans* (18–20). To test our hypothesis, we ran the square barrier experiment with adult Day 7 nematodes. In contrast to young adults, only 41% of Day 7 *C. elegans* have crossed the barrier after 120min (Fig. 6C, purple diamonds). When adult Day 7 nematodes were tested without a physical barrier (control experiment), results were similar to the ones of adult Day 1 nematodes (Fig. 6C, pink diamonds). Since Day 7 adults are beyond middle age, the above findings suggest an aging-related change in *C. elegans* physical barrier crossing behavior.

### The effect of confined area size

Next, we asked whether the size of the confined area and its ratio over the total available plate area affects the dynamics of the assay, i.e., whether worms travel in and out of the target area or stay inside the target square once they cross it.

To this end, we ran the same experiment, using a 5×5mm 3D-printed NGM square instead of a 20×20mm one. Indeed, when the 20×20mm square was used (Fig. 6), the nematodes that crossed the barrier stayed inside the framed area for the remainder of the assay. In case of the 5×5mm square, (Fig. 7C *vs* Fig. 6B), nematodes did not remain inside the square once they enter, instead they might enter, roam, exit, and even reenter. Due to this dynamic situation, the number of worms counted inside the square did not increase monotonically, as in Fig. 6, instead, it fluctuated. For this reason, and to reflect this dynamic behavior, instead of a time course (Fig. 6C), results are presented as scatter plots (Fig. 7A, 7B), showing the mean and standard deviation of the percentage of worms scored inside the framed area at any given time point.

In addition, the 20×20mm square (Fig. 6) frames ∼15% of the 60mm plate surface area, whereas the 5×5mm one (Fig. 7) frames <1% of that area. This is probably why a smaller percentage of worms is scored inside the 5×5mm square at any given moment, compared to what happens with the 20×20mm one. Therefore, the relative size of the square with respect to the culture plate affects the dynamics of the assay and should be taken into consideration when data is interpreted.

### The effect of feeding history

When two groups of Day 1 adults were tested (Fig. 7A), a fully fed one (FF; purple inverted triangles) and one that was starved for 24 hrs prior testing (S; blue diamonds) it was found that starved animals enter the food-baited square in higher numbers than fully fed ones. This suggests that starved animals might have a stronger motivation to explore their surrounding area and possibly to overcome physical obstacles, as well.

When the experiment was performed with no OP50 inside the square (Fig. 7A, woF-without food, shaded area, *vs* wF-with food, non-shaded area) *C. elegans* behaved similarly, and starved animals (S woF, blue circles) entered the food-baited square in higher numbers than fully fed ones (FF woF, purple triangles). Therefore, the nematodes’ feeding history has a strong influence on their tendency to explore beyond a physical barrier even if there is no food beyond it.

### The effect of prior experience

Lab populations of *C. elegans* are commonly grown either in liquid cultures or on flat NGM plates with practically 2-dimensional surfaces. *C. elegans* used in the present work have been cultured for many generations on NGM plates. Hence, we asked whether a group of nematodes that has dwelled on a NGM plate featuring a 3D-printed square will have familiarized themselves with it, and thus will cross a similar 3D barrier in higher numbers. To this end, we placed L4 nematodes on a seeded NGM plate with a 5×5mm Parnon-printed square, for 24 hours. Then, and on Day 1 of their adult life, we tested them on a different plate, equipped with a similar, food-baited 5×5mm square.

Nematodes that have previously experienced an environment which includes a 3D square (Fig. 7B, 3D), cross the barrier in higher rates than the ones that have experienced only the flat surface of a regular NGM plate (R). This is the case regardless of whether the 3D structure frames a food baited area (wF) or a non-baited area (woF, shaded).

### Spatial restriction of egg laying

We explored whether the Parnon-printed squares can be used to spatially control selected *C. elegans* behaviors, i.e., egg laying. To this end, a 5×5mm Parnon-printed square, baited with OP50, was used. (Fig. 8). We used the small size squares to apply a stricter constrain on the behavior we aim to control. Results showed that Day 1 adults that were placed on the plate and were left there for 24 hours, laid eggs almost exclusively in the confined area or on the square barriers themselves, since almost no eggs are found at other plate locations (Fig. 8A-D).

**Figure 8:**
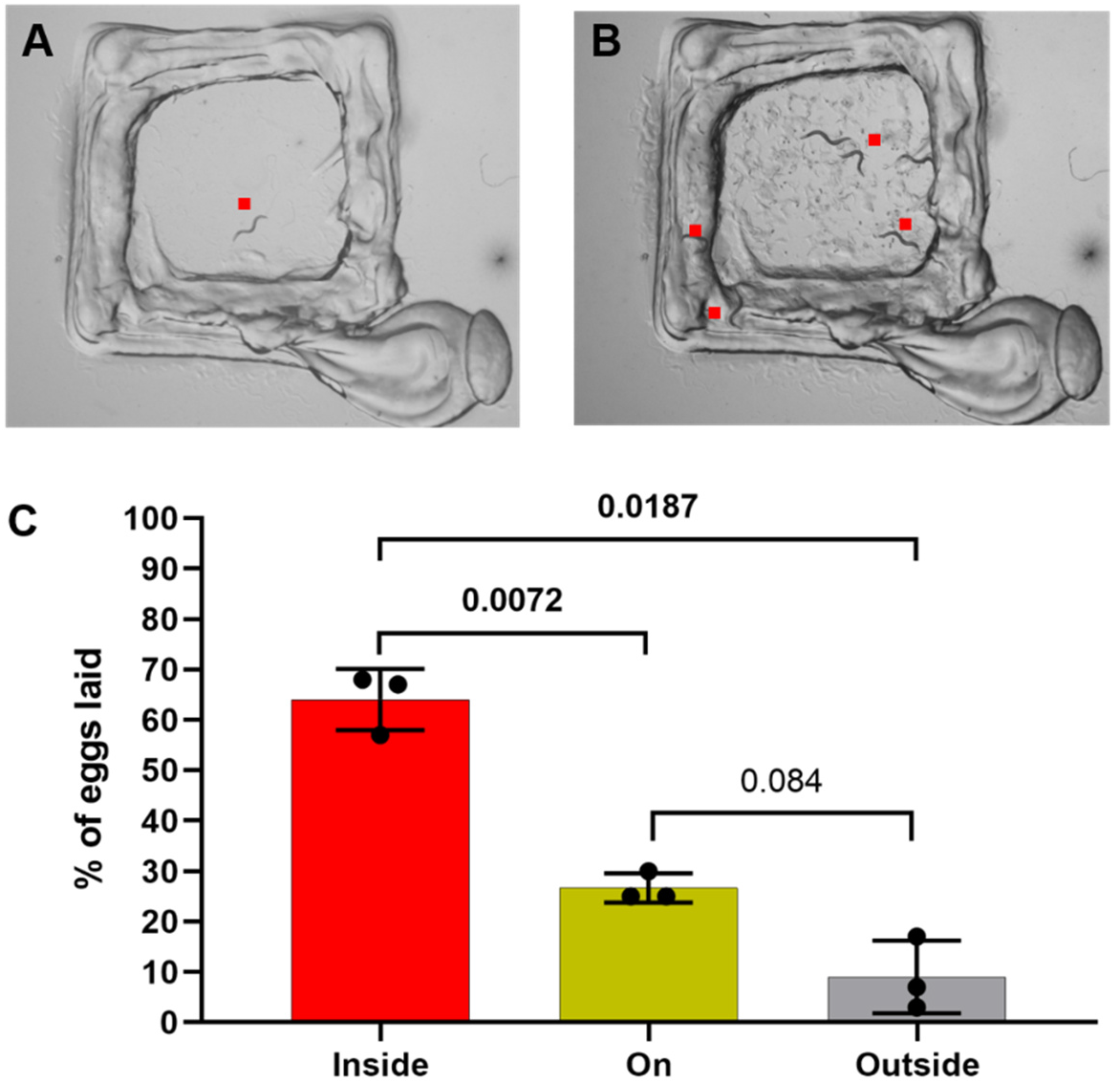
3D-printed structures can be used to confine *C. elegans* behavior. (**A**) 3D-printed square (5×5mm, 3 layers, 0.5mm thick each), seeded with OP50, on an NGM plate, at time t=0. (**B**) The same square as in A, after 24 hours (t=24hr). Eggs have been laid only inside the confined area, and several nematodes can be seen on and inside the square. Note the absence of eggs in the proximity of the square frame. Nematodes initially placed as in Fig. 7C. (**A**) and (**B**): red dots indicate location of *C. elegans*. (**C**) Graph showing the % of eggs laid in the region of each 3D square tested, as shown in A and B, inside: eggs laid inside the framed area, on: eggs laid on the square frame, outside: eggs laid in the premises of the squares, in 5mm distance. Bars: mean, error bars: SD, dots: 3 individual experiments (worms used in each experiment: n_1_=10, n_2_=12, n_3_=11), above bars: p-values of indicated comparisons, student’s paired t-test, with bold p-values<0.05 (significant difference).

## Discussion

Less than a handful of attempts has been made to date to establish worm-friendly and experimentally informative 3-dimensional arenas for *C. elegans* (21–23). In two of these studies, the 3-dimensional platforms are made of porous materials, primarily intended for cultivating and imaging worms (21, 23). In the third case (22), nematodes swim in microfluidic devices that resemble their granulated natural environment, i.e. soil. These substrates provide a significantly more realistic terrain than the NGM plate, however each of them lacks one of the key traits (well-defined and structured arenas, compatibility with microscopy and imaging techniques, or easy and cheap fabrication) that would allow it to be widely adopted for 3-dimensional behavioral experiments. The methods presented here, especially the Parnon NGM printing, aspire to fill this gap.

### PVA-casted 3dimensional structures

PVA casting successfully results in creating diverse 3-dimensional NGM structures (Fig. 2). However, there are important downsides. When the chambers of the PVA cast (Fig. 2A) are small (∼1mm in diameter), NGM is not casted properly. This could be related to the fact that small spaces result in faster cooling of NGM, and subsequent clogging. A possible way to resolve this would be to keep the cast warm throughout the process to prevent premature NGM solidification.

At the same time, some of the cast channels might be partially blocked, due to trapped air bubbles or 3D printing discrepancies, occasionally happening when such small empty spaces need to be achieved. A 3D printer with higher resolution than the Ultimaker3 could potentially help resolve this issue, increasing, however, the overall cost of production.

Lastly, PVA casting results in parts with very rough surfaces (Fig. 3). The increased roughness of the NGM surfaces may or may not be a desired feature, depending on the experiment to be conducted. Indeed, strong roughness could possibly impede or redirect nematode locomotion or interfere with optics and imaging. Note, for example, how challenging is to distinguish a nematode that is crawling on the NGM casted structures of Fig 3 (3A: yellow arrow, 3B, 3C: yellow boxes).

In addition, PVA casting process is time consuming, as it takes ∼28 hours per part to complete (see Methods). It is also resource heavy, since PVA is significantly more expensive than most PLA or other standard plastic filaments. Moreover, the casts are not reusable, so a new cast must be printed and dissolved for each new NGM part. Furthermore, access to a high-resolution printer, like the Ultimaker3, is required. In addition, certain features cannot be obtained using casts, as for example complex vertical elements, sharp angles, and very small size features. Consequently, PVA casts do not constitute a go-to option. This outcome prompted us to explore the 3D-printing route discussed below.

### Parnon customization and Parnon-printed 3D-structures

The printability of hydrogels has been broadly demonstrated (15, 24–26), including the extrusion of agarose-based hydrogels (27, 28). The 2% agar concentration in the NGM mix is among the lowest used in hydrogel 3D printing (17, 26, 29) without the use of enhancers (30). Moreover, the Parnon-produced structures presented here and used in *C. elegans* behavioral experiments feature three layers of NGM. This constitutes a promising achievement when compared to previously reported lattices and scaffolds, made with hydrogel ink of similar concentration and properties (15), mainly because it was possible without additional support (31). In coarse trials where multi-layer cylinders or walls were attempted, although the stability of the structures was satisfying, the overall printing quality was low and inconsistent (Supplementary Fig. S9).

NGM lines can be laid successfully in 90° angles or smaller (Suppl. Fig. 7B). This allows for a diverse set of designs, although the printing quality is compromised (Supplementary Fig. S7B, yellow frame). Printing lines in sharp angles is in general a challenging task (26) and the performance of Parnon is currently considered satisfactory for the current experimental needs. Stricter control of the extrusion process would improve this aspect (26).

In general, the resolution of the FDM-printed structures is restricted by the nozzle diameter (32), which is limited by clogging, and depends on the rheological properties of the extruded material. Hence, the typical resolution for FDM is ∼100μm (32). When it comes to bioprinting hydrogel inks that contain biomolecules, the typical size of the printed features is ∼500μm (32). According to other reports (33), the achieved resolution in non-hydrogel specific FDM printing is around 50/250 μm (*z/xy*), while older actuation pressure extrusion attempts using hydrogels (34) report a *z/xy* resolution of 500 μm. The resolution achieved by Parnon reaches 300μm in *z*, and ∼250μm in *xy*. Therefore, Parnon’s performs well regarding its *xy* resolution, when compared to overall FDM printing, and scores better than other hydrogel-ink printers. Parnon’s *z* resolution ranks close to its hydrogel-ink peers but is considerably worse than generally achieved FDM *z* resolution. Better control of the actuation pressure and extruded NGM viscosity could help improve this property. Given the size of adult *C. elegans’* body, i.e. ∼1mm length and 70-100μm diameter (35), a moderate upgrade would probably suffice for most applications. Nevertheless, the resolution achieved by the current version of Parnon serves well the experimental purpose of the printed parts.

Compared to the PVA-casted parts, Parnon-printed structures are smoother (Fig. 5 *vs* Fig. 3), allowing for easier imaging of nematodes (Fig. 5B inset). Apart from the initial purchasing cost of the commercial printer, 3D-printing hydrogel arenas using Parnon is an affordable and fast way to create 3-dimensional assays, especially when compared to the PVA casting.

### Parnon mechanics, extrusion, and software communication

Currently, pausing and resuming printing is a challenging process. Unlike market FDM 3D printers, the extrusion of which can start and stop easily on demand, Parnon still struggles to do that quickly and precisely. The result is undesired dripping of NGM, even when the actuation pauses or reverses. Because of this, the printing process needs to be continuous (Supplementary Fig. 7), thus affecting the printing paths of certain designs (e.g., cross shape, Supplementary Fig. 7C). Deeper understanding of the compressive behavior and solidification kinetics of NGM, more efficient control of the nozzle heating, or interventions at the extrusion pathway could result in more precise extrusion control and ultimately allow for more tailored designs.

Parnon’s custom print head is not compatible with the basis printer firmware and thus requires its own commands. Market available gCode slicers do not have the capability to output relevant and processable linear actuator commands in the Parnon’s format. The development of the gCode and Arduino commands remains a rigorous manual process. We are working toward implementing appropriate gCode slicers in future iterations of the prototype.

### C. elegans ability to cross physical barriers

To demonstrate the suitability of the Parnon-printed parts as *C. elegans* behavioral arenas, we showed that three-layered NGM squares can be successfully used to explore nematodes’ ability to cross physical barriers (Fig. 6). This is the first attempt to explore *C. elegans* ability to overcome physical obstacles, made of the same nematode-friendly material on which worms are cultured in the lab.

The height of the square used (Fig. 6), is estimated at ∼1.5mm (three layers, 0.5mm thick each), which equals approximately 1.5times the adult *C. elegans’* body length. This means that the hurdle is not extraordinarily high, compared to *C. elegans*, yet it is hard to establish a reference, since similar experiments have not been conducted before. Having in mind the 3-dimensional character of *C. elegans*’ natural habitats (5, 36), we speculate that an environment with 1.5mm high features is not out of range for a field-dwelling nematode. Therefore, we assume that such encounters are not uncommon for *C. elegans* in the field. Hence, the finding that young adult worms can easily climb over them is not surprising. It is noted that in addition to nematodes which climb over the square barrier, ∼5% of the population tested crawled between the square and the substrate on which it rested.

Basic motor skills and chemotaxis remain functional through later stages of adulthood in *C. elegans* nematodes (37), however some decline in locomotion speed (38) and spontaneous locomotion has been reported (19, 39). The observed difference between Day 1 and Day 7 adults (Fig. 6) could be attributed to age-affected locomotion performance.

Results from dispersal assays have shown that young adult *C. elegans* move away from the original position and spread into a wide area on the assay plate, but older adults are more likely to remain close to their original location (19). This agrees with our findings on Day 7 adult nematodes, which do not spread over the square barrier as their Day 1 conspecifics (Fig. 6). Dispersal over physical obstacles has not been previously assessed. Notably, a non-negligible part of the Day 7 population in study did cross the barrier, revealing a diversity in individual *C. elegans* behavior (40) and maybe a diverse impact of aging-related changes (41).

It is reported that learning and memory decline earlier in adulthood than other physiological operations (7, 37, 42) and show signs of deterioration on Day 7 (37) or even Day 5 of *C. elegans* adult life (7). It is possible that some aspects of decision making, e.g., related to overcoming physical challenges, are affected, as well. Moreover, a decrease in food-seeking exploration in older nematodes is possible.

Our findings on starved *C. elegans* (Fig. 7A), are in alignment with known starvation-induced modifications of nematode behavior that improve their food location likelihood. Indeed, after prolonged starvation, *C. elegans* change their strategy to long-range dispersal (43–46). Even after 10–20 min away from food, neuronal plasticity can change, leading to global, instead of local, search behavior in adult nematodes (47). Moreover, it has been shown that food deprivation increases threat tolerance in *C. elegans*, resulting in worms that make “bolder” decisions (48). Broadening the explored area, even if it includes some physical challenges, may be an additional manifestation of behavioral changes triggered by food deprivation. Feeding history seems to impact obstacle crossing more than the presence or absence of food inside the framed area (Fig. 7B).

*C. elegans* nematodes used in the present work have been cultured for many generations on NGM plates, conforming with common practice in the field. Consequently, the 3D-printed squares are practically the first 3-dimensional structure they encounter, besides the unfriendly walls of the plastic petri dish itself and the NGM chunk often used in culture maintenance, when transferring nematodes into a fresh culture plate. In case of the latter, *C. elegans* usually actively interact with the chunk, but they commonly just crawl away from it, since the chunk is flipped over the fresh NGM plate (6). Our findings (Fig. 7B) suggest that *C. elegans* behavior with regards to physical challenges is affected by their prior experience and familiarity with similar structures. We expect that this effect will be stronger if the animals are grown in a 3-dimensional environment since hatching, and more 3-dimensional structures exist in their culture plate.

### Spatial confinement of C. elegans behavior

Given that in the above experiments (Fig. 7) *C. elegans* get in and out of the square frames without much difficulty, the localized egg laying behavior (Fig. 8) can be attributed to the fact that the only food source available is located inside the 3D squares. This makes them a preferable location for egg laying since this way the newly hatched progeny has immediate access to food. Indeed, it is known that egg laying is regulated by a number of environmental conditions (49), leading to spatial control of the egg laying behavior. Moreover, the egg laying rate in the presence of abundant food is significantly higher than in the absence of food (50). The use of the 3D squares works well with localized food availability and it proves very effective when it comes to restricting egg laying, and potentially the hatching of L1 larvae, as well.

## Conclusions

This is the first reported attempt to explore *C. elegans* ability to cross physical obstacles, made of the same nematode-friendly material on which worms are cultured in the lab. A method that enables fabrication of such structures, as is the prototype NGM-extruding Parnon 3D printer, was the necessary condition to achieve this. We demonstrate that new tools and assays are not only the key to answering biological questions but, most importantly, can often shape the very questions posed. We envision the applications of 3D printing NGM to enable artificial landscaping and development of diverse playgrounds for the exploration of *C. elegans* 3-dimensional behavior.

## Methods

In this section we describe the PVA casting and the Parnon printing methods, and the *C. elegans* behavioral experiments process. Technical details on the printer’s customization, NGM rheological properties, and software communication between the printer’s parts can be found in the Supplementary Information section. A list of the major components used for the conversion of the commercial printer into Parnon us provided in the Appendix. Details on the Arduino code are provided in the Appendix.

The material used to build 3D-behavioral arenas is NGM (Nematode Growth Medium), the agar-based hydrogel used to culture *C. elegans* in the lab. NGM 2% in agar was prepared according to standard methods (6, 51). Once all the ingredients are diluted in water, NGM is in liquid form in temperatures higher than ∼45°C (melting point), and it solidifies as it cools down. In room temperature, polymerized NGM is solid. This is a key property for both the PVA casting and Parnon 3D printing methods.

### PVA casting-PolyVinylAlcohol; PVOH; [CH2CH(OH)]n)

This approach takes advantage of the water solubility properties of PVA. To design the cast, SolidWorks (Dassault Systemes, France) was used. The cast was printed at an Ultimaker3 3D printer (Ultimaker BV, United States). The casts shown in Fig. 2 were printed in ∼5 hours. Once the cast was ready, liquid NGM was injected in. NGM took the shape of the space that is left vacant inside the cast. Once the NGM solidified thoroughly, after ∼30 min, the cast was immersed in deionized water and was sonicated for ∼24 hr in a sonicator bath (Branson Ultrasonics, USA). This accelerated dissolving of the PVA cast, which was entirely dissolved and was therefore not reusable. Since NGM is not water soluble after polymerization, at the end of this process only the NGM structure was left intact.

### Fused Deposition Modeling (FDM) 3D printer frame

We used a commercially available FDM printer, the DOBOT MOOZ-2 machine (Shenzhen, China, Fig. 4). It was preferred against other options due to i) existing gCode interpretation firmware that controls the linear actuators in the cartesian axis system, ii) price efficiency, as a result of low build volume, and iii) frame rigidity. According to the manufacturer, MOOZ-2 is made of aircraft grade aluminum alloy, which minimizes *in situ* vibrations and increases rigor.

All three axes are controlled with lead screw linear actuators which are more accurate, precise, repeatable and user-friendly than the alternative. The MOOZ-2 is amenable to modifications (Fig. 4, Supplementary Fig. S1) that allow Parnon’s custom print head and substrate to be attached.

### Parnon custom substrate

Parnon’s substrate is designed to limit liquid spreading and promote build up along the z-axis (Fig. 4). The custom substrate consists of three major parts: i) the plotting medium (Fig. 4 Part Di & ii), ii) the cooling apparatus (Fig. 4 Part Diii & iv), and iii) the leveling system (Fig. 4 Part Dv).

#### i. Plotting medium

We use a plotting medium (34, 52, 53) to reduce liquid spreading, provide support and promote the cooling process. The plotting medium (Fig. 4 Part Dii) is 1.4% glycerin (Sigma Aldrich, USA) in water. It has the same density as liquid NGM (1.024 g/mL), so the extruded NGM is effectively suspended during the print to assist in limiting liquid spreading. A 60mm diameter glass petri dish (Fig. 4 Part Di), holds ∼20mL of the plotting medium for each printing session.

#### ii. Cooling apparatus

NGM is heated by the heating element when inside the printing head (Fig. 4C, Supplementary Figure S3A). Moreover, there is Joule heating around the nozzle, to keep the NGM liquid during extrusion (Supplementary Figure S3). To cool down the plotting medium and facilitate NGM solidification, a Peltier device (Northbear Electronics) is used (26) (Fig. 4 Part Diii), operating in 16V 12A. This device cools the plotting medium according to the thermoelectric effect (Fig. 4 Part D, red arrows). An aluminum heat sink (Fig. 4 Part Div) is implemented underneath the Peltier device to dissipate heat and ultimately increase its efficiency. The Parnon’s cooling mechanism operates at 86% efficiency, achieving a 2.0°C temperature decrease from the surface of the plotting medium to the floor after ∼20min (Supplementary Fig. S6). Details on the efficiency of the Peltier device and the heat flux generated are provided in the Supplementary Information.

#### iii. Leveling system

The Parnon uses four individually adjustable spring resistance screws (Fig. 4 Part Div, Dv) to allow for substrate leveling. Additional circular levels with adjustable screws are placed on the petri dish (Fig. 4 Part Di, Supplementary Fig. S2).

### Parnon custom print head

The custom print head consists of three major parts: i) a mechanism to house and heat NGM (Fig. 2 Part Civ, v, vii, viii, ix & x), ii) gCode and Arduino software communication, and iii) a mechanism to provide actuation pressure (Fig. 2 Part Ci & iii).

#### i. Housing and heating mechanism

The aluminum heat sink (Fig. 2 Part Cv) was machined from 6020 aluminum. It connects all the components on the custom print head to Parnon’s x axis. The heat sink is heated with a 3” 3/8” diameter 200W 120V heating element with an internal k-type thermocouple (Fig. 4 Part Civ), which is controlled by a programmable temperature controller. The temperature controller is set at 65 °C because we expect G’=∼5Pa (Supplementary Fig. S6). This modulus is sufficiently low to allow easy extrusion and sufficiently high to limit liquid spreading. Copper wire with 5V potential (Fig 4. Part Cx, Supplementary Fig. S3) heats the nozzle (joule heating effect) so NGM does not solidify while in the nozzle.

The current version of Parnon uses a 3mL glass syringe (Fig. 4 Part Cviii, Supplementary Fig. S3) to house liquid NGM for extrusion. An early version used a 5mL glass syringe. The glass plunger (Fig. 4 Part Cvii) is connected to the linear actuator arm (Fig 4. Part Ciii) via a custom 3D-printed connector (Fig. 4 Part Cvi, Supplementary Fig. S3, Formlabs Tough Resin), designed using Solidworks (Dessault Systemes, France).

NGM is extruded through a ½” stainless-steel luer-lock nozzle. Various gauges of luer-lock nozzles were tested (higher gauge translates in lower inner diameter). The initial nozzle had a 813 *μm* ID, we improved to a 404 *μm* ID nozzle, and the current version features a 254 *μm* ID nozzle. Print resolution improves as the ID of the nozzle decreases (Fig. 5, Supplementary Fig. S7).

#### ii. gCode and software communication

Modifying the hardware of an existing 3D printer compromised the communication the printer had with the original print head. To resolve this issue, we introduced a limit switch, an Arduino card, and a stepper motor driver. More details are provided in the Supplementary Information section.

#### iii. Actuation pressure

A linear actuator facilitates extrusion by compressing NGM in the syringe. A characteristic stress is required for the NGM to begin flowing through the nozzle. The time required to reach stress (*t_eq._*) is the target of our analysis of NGM under actuation pressure. Time *t_eq_*. is used to guide the amount of time prior to print extrusion beginning dedicated to reaching the characteristic stress (strain). More details are provided in the Supplementary Information section.

### NGM material characterization

NGM 2% agar was prepared according to standard methods (6, 51). The density of liquid NGM was experimentally evaluated to be ∼1.024g/mL ( ∼10mL of NGM weight ∼10.24g, and the density of deionized water was considered 1.0g/mL). The melting point of agarose (molecular biology grade, Sigma Aldrich, USA) is ≤65°C and the transition temperature (gel point) is 36°C ±1.5 °C (for 1.5% gel), according to the manufacturer. 100microliters of blue food color (Americolor, CA, USA) were added to 100mL of NGM (0.1%) for *in situ* visibility and observational purposes.

#### i. NGM rheology

Rheology experiments were performed on NGM to uncover the fastest solidification temperature, using a TH DR2 Rheometer (TA Instruments, USA). The temperature was dropped from 60°C to 25°C at a rate of 5°C/min. The optimal solidification temperature is determined by the maximum slope of the G’ *vs* T curve (Supplementary Fig. S6). The slope (dG’/dT) peaks at 35.9°C (Supplementary Fig. S6), indicating the fastest solidification temperature.

#### ii. NGM compressive viscoelastic response

NGM presents a viscoelastic response to compressive stress. Compressive stress tests were run on NGM at 65°C in a 9.11mm ID (inner diameter) and a 12mm ID glass syringe. Time *t_eq_*. is required to reach the inflection point of the extrusion equilibrium stress (*σ_eq_*.) under actuation pressure, and time *t_r_* is required to relax from it. The inflection point occurs at a stress value *σ_eq_* which varies with strain rate *ε̇*. The time *t_eq_*. required to reach it, varies depending on syringe ID.

Details on NGM viscoelastic response characterization are provided in the Supplementary Information section.

### C. elegans behavioral experiments

Snapshots of *C. elegans*’ actions during behavioral experiments were taken with a DP22 camera, mounted on a SZ61 dissection microscope, using CellSens Software (all by Olympus, Japan).

#### i. Barrier crossing

In the control experiment, where *C. elegans* are allowed to reach a food source not framed by a physical barrier, a droplet of OP50 was placed on a regular NGM plate, and was framed by a figurative square, drawn with a marker on the bottom of the plate. Nematodes were placed around the targeted area (Fig. 6D, left) and were allowed to roam free.

In all experiments where Parnon-printed squares were used (Figs 6, 7 and 8), the 3D structures were rinsed multiple times with deionized water and then were placed on a 60mm NGM plate. Next, when applicable, a of *E. coli* OP50 was pipetted gradually inside the square and was left to dry for ∼10min (10microliters for Fig. 6 experiments, 5 microliters for Figs. 7 and 8 experiments). A number of *C. elegans* nematodes were then transferred on the NGM plate, outside of the square(s), at least 3mm away, at random locations.

To explore the effect of aging on crossing physical barriers (Fig. 6), two age cohorts of adult hermaphrodites were tested, namely young adults of Day 1 (L4+1), and middle-aged animals of Day 7 (L4+7). We used a ∼20×20mm 3D-printed square, made of 3 NGM layers, 0.5mm thick each (see Methods). We considered t_0_ the time point at which all nematodes were placed on the plate. The worms were allowed 120min to explore, and worms scored inside the square were counted every 5 or 10min. Nematodes that were on the square were counted towards successful crossings. The *E. coli* OP50 food used in all trials came from the same stock batch, and experiments were run on 3 different days.

To explore the effect of feeding history (Fig 7A, 7B) we tested young adults of Day 1 (L4+1), and middle-aged animals of Day 7 (L4+7), that had either been fully fed (FF) or starved for 24hrs (S). To explore the effect of prior experience (Fig. 7C) we tested young adults of Day 1 (L4+1) that had been either grown on a regular NGM plate (R) or had been moved into an NGM plate containing a Parnon-printed square, similar to the one used in testing (3D). In all cases (Fig. 7A, 7B, 7C) both a baited (wF) and a non-baited 3D square (woF) were tested. Moreover, in all cases (Fig. 7A, 7B, 7C) we used ∼5×5mm squares, consisting of 3 layers 0.5mm thick each.

#### ii. Spatial control of egg laying behavior

For these experiments (Fig. 8), ∼5×5mm3D-printed NGM squares consisting of 3 layers 0.5mm thick each, were used. After printing, the squares were rinsed with deionized water, and then were placed on a 60mm NGM plate. Next, 5microlitters of *E. coli* OP50 were gradually pipetted inside the square area and were left to dry for ∼10min. A population of Day 1 (L4+1) adult *C. elegans* hermaphrodites was then transferred on the NGM plate and were placed at least 3mm away from the squares. The plate was checked for eggs after 24hrs.

### C. elegans strains

N2 Bristol (wild type) *C. elegans* were used in all experiments, initially acquired from CGC (Caenorhabditis Genetics Center, provided by the *C. elegans* Reverse Genetics Core Facility at the University of British Columbia, which is part of the international *C. elegans* Gene Knockout Consortium) and maintained in the lab.

### Statistical analyses

Statistical evaluations were made using t-tests (GraphPad Prism 9.0.0) with *p* values considered significant when *p*>0.05. In Figure 6, multiple unpaired t-tests were performed, and in Figure 7 evaluations were made using two-tailed, unpaired t-tests. Additional information is provided in the figures’ captions.

## Acknowledgements

We thank Ao-Lin Hsu for the use of space and equipment, and Nikos Chronis for the use of space. We are grateful to Zijun (Justin) Yuan for hydrogel testing, to Chris Pannier for technical advice, to Bill Kirkpatrick and Kent Pruss for assistance with machining, to Alex Shorter for use of Ultimaker3 3Dprinter, to Michael Solomon and Yufei Wei for rheometry advice, to Arthur Sinclair for rheometer facilitation, and to Ruiming Lu for help with glass cutting. This work was funded by the University of Michigan Office of Research (UMOR)-Faculty Grants & Awards Program (EG). EG is the recipient of a NIH-NIA K01 award. The content is solely the responsibility of the authors and does not necessarily represent the official views of the National Institutes of Health.

## Authors’ Contributions

SNC designed and printed the PVA molds, built the Parnon printer, performed analyses, collected data, generated figures, and wrote the manuscript. LYLT assisted with customizing the commercial printer, and reviewed manuscript. KB contributed advice and expertise, and reviewed manuscript. EG conceived the idea, designed Parnon experiments, designed and performed *C. elegans* experiments, analyzed data, generated figures, wrote, and edited the manuscript, and supervised research.

## Supplementary Figures and Tables

**Supplementary Figure S1:**
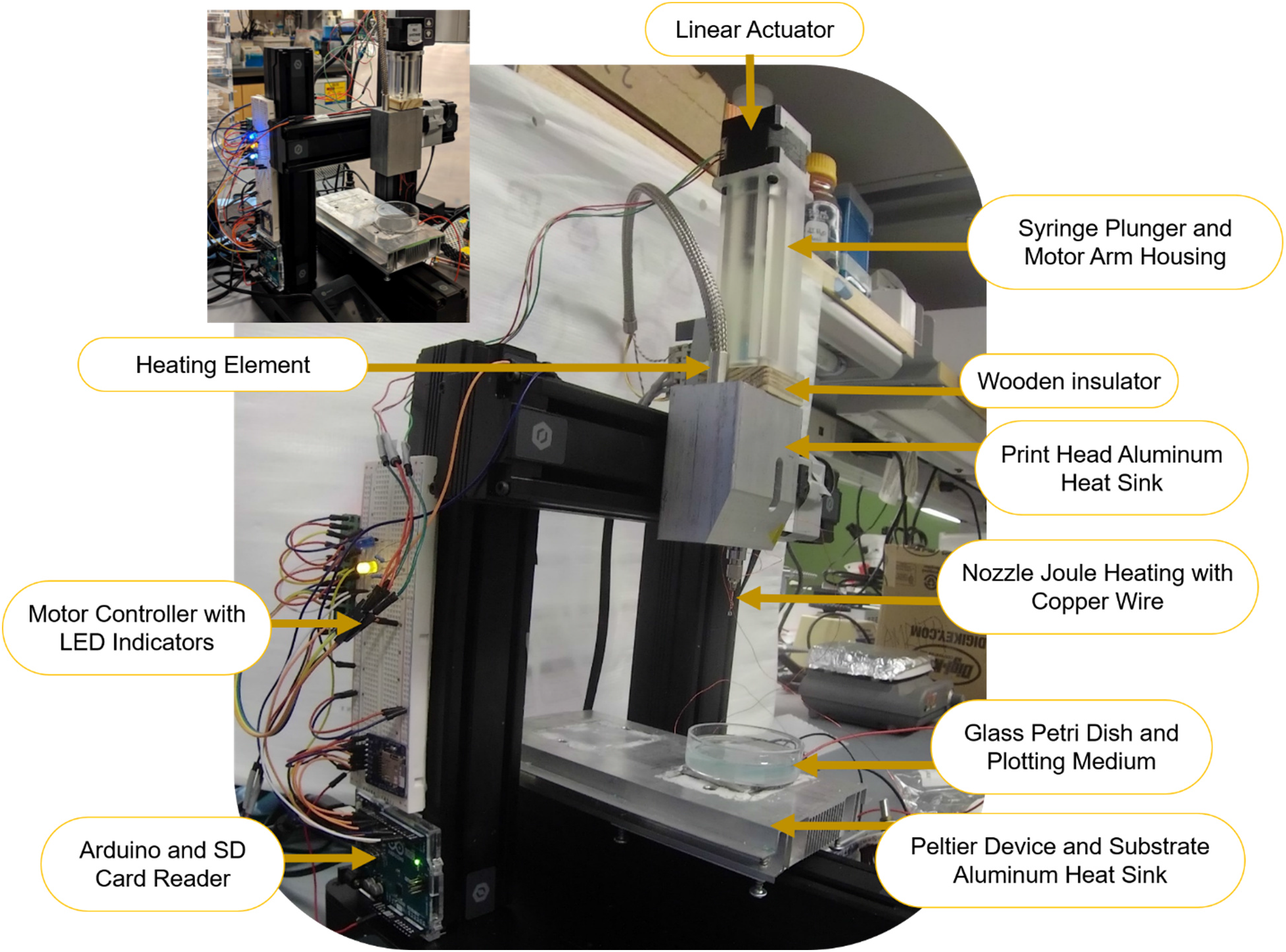
Overview of the Parnon printer. The major customized and house-made parts are indicated. Small inset: early version of the Parnon printer, in which a shorter linear actuator and syringe housing are featured.

**Supplementary Figure S2:**
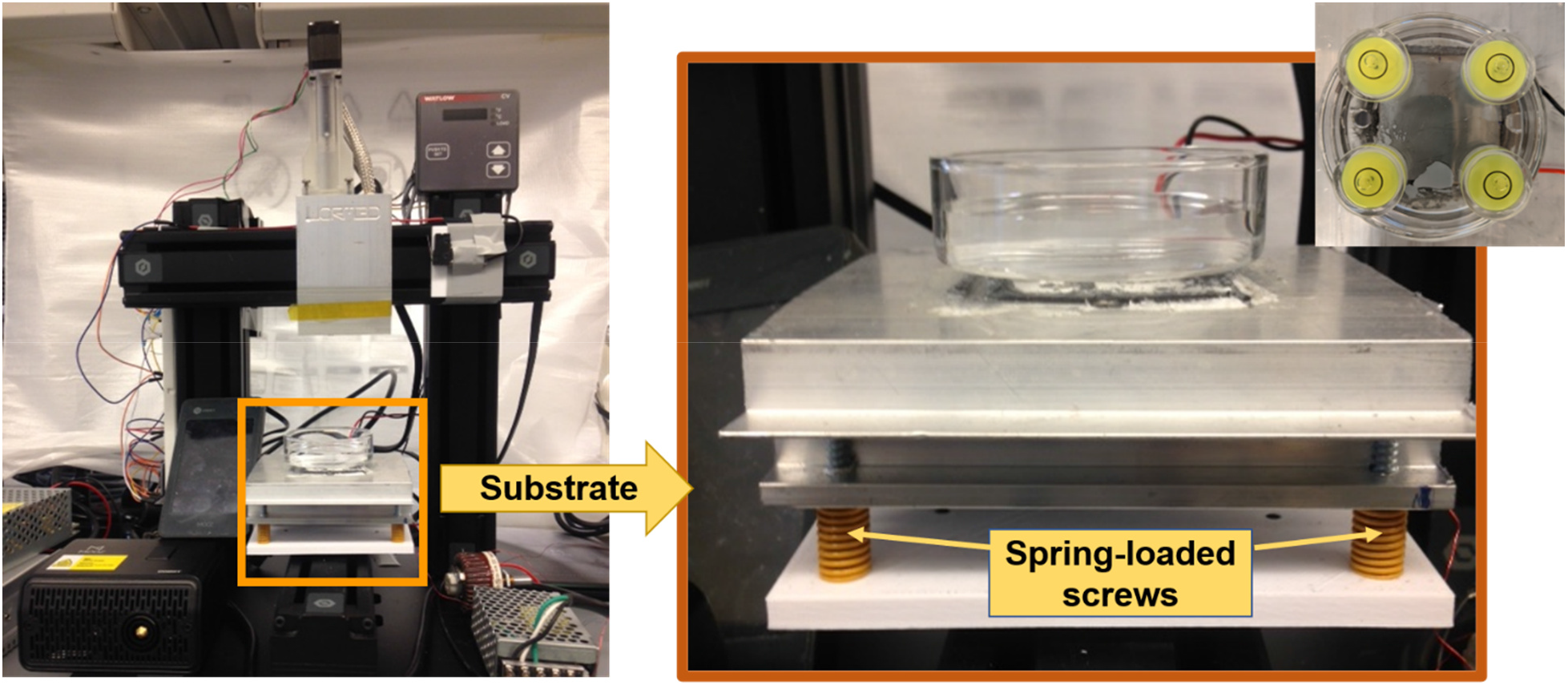
Four-point bed leveling system. Each corner of the substrate is equipped with a spring-loaded screw, to either raise or lower the corresponding corner of the substrate. Circular levels are placed at each corner of the substrate (small inset) and the screws are adjusted until each level reading matches the rest.

**Supplementary Figure S3:**
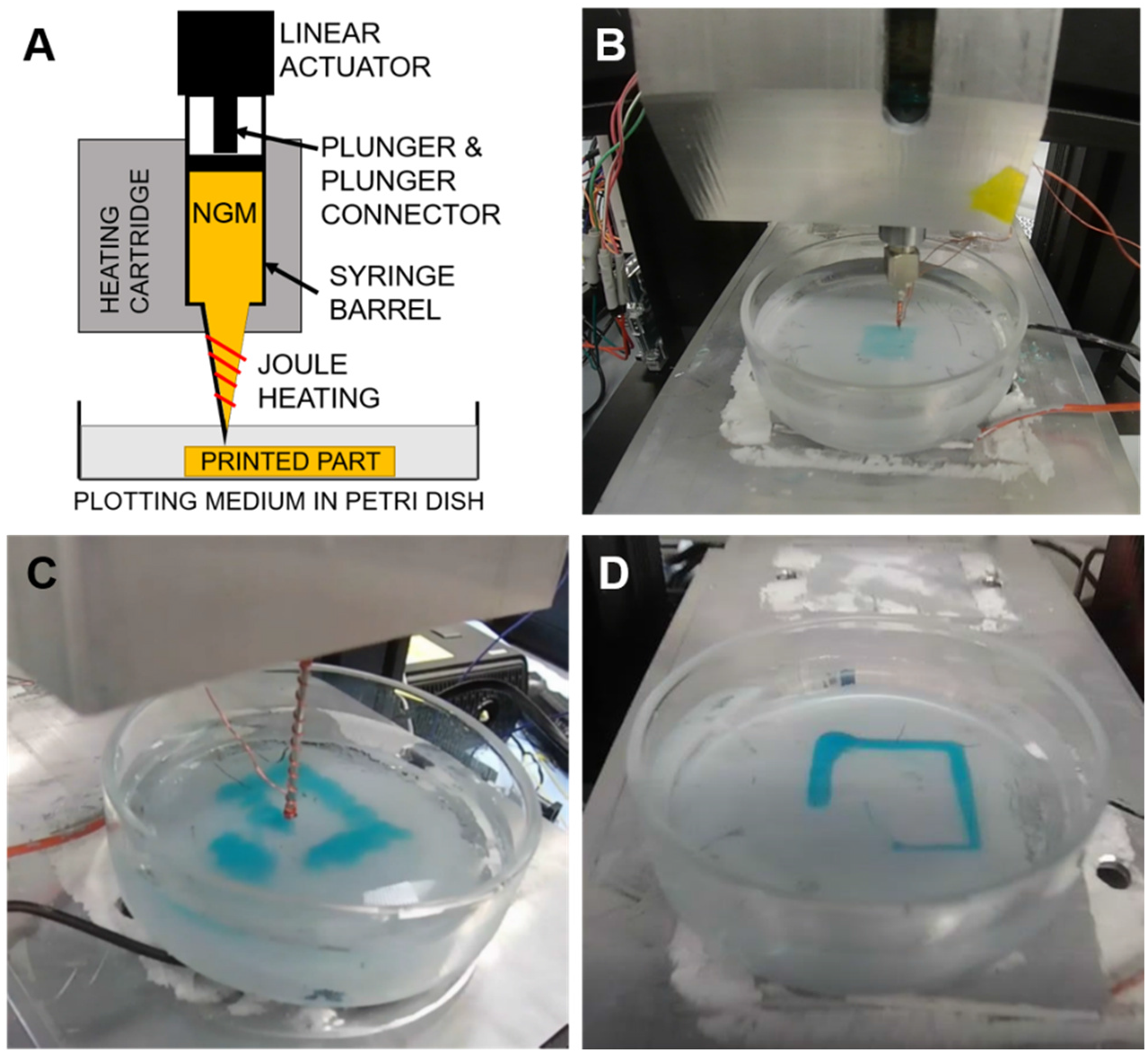
The printing apparatus and the nozzle joule heating detail. (**A**) Schematic of the syringe barrel containing liquid NGM, sitting inside the heating sink which contains the heating cartridge. The joule heating is illustrated around the protruding nozzle. (**B**) Printing a two-layer pad, as shown in Suppl. Fig. S3 A&B. (**C**) Printing a square structure; note the wider nozzle which results in thicker printed lines, compared to (D). A longer nozzle is used for demonstration purposes. (**D**) A semi-finished square, still in the plotting medium; note the thinner lines compared to (C), as a result of the narrower nozzle used.

**Supplementary Figure S4:**
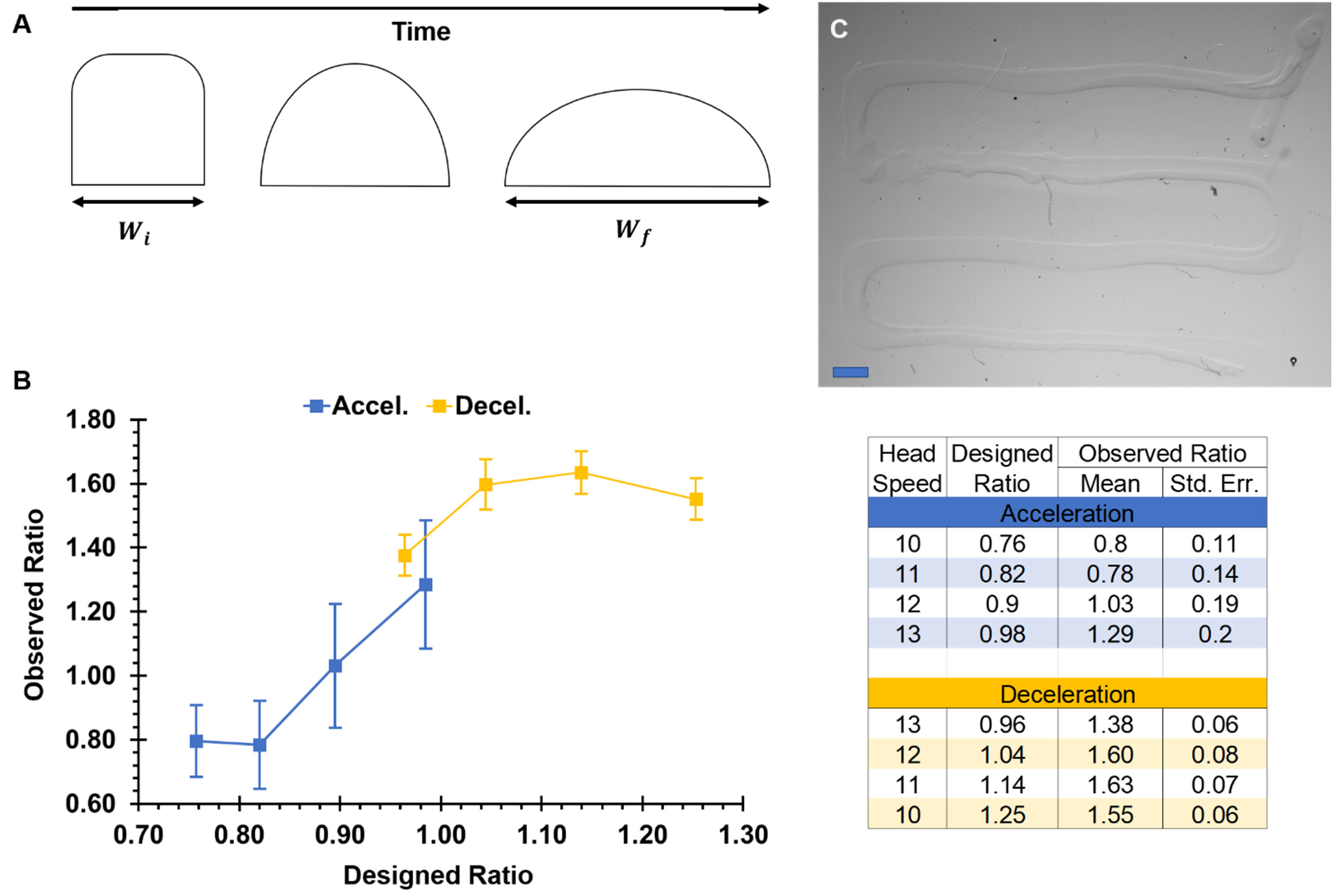
Liquid spreading control and line width experiments. (**A**) Schematic of the cross section of a printed line that exhibits liquid spreading, evolving over time. The initial width (*W_i_*) is the line width immediately after the line is extruded, which is approximately equal to the width of the nozzle. As time passes, the width of the line increases while the height decreases. The final width (*W_f_*) is the line width after the equilibrium shape has been reached. (**B**) Graph showing the relationship between the ratios of designed line widths and observed line widths to *W_i_*. A designed ratio of 1 means the expected value for *W_f_* is equal to the width of the nozzle. The designed line width is the expected value for *W_f_* based on user defined extrusion volume. An observed ratio of 1 means the final width (*W_f_*) is equal to the width of the nozzle. The observed line width is the measured value for *W_f_* after printing. Means with standard error bars are shown. To achieve an observed ratio of 1 (*W_f_* = nozzle width) the design ratio needs to be ∼0.90. Table shows data from line width experiments. (**C**) Image from the line width experiments. Four consecutive 20 mm long lines were printed with a head speed ranging from 10 *mm/s* to 13 *mm/s*. In four experiments, the head speed started at 10 mm/s and incremented 1 *mm/s* per line with an actuation speed of 13.4 *μm/s*. In another four experiments, the head speed started at 13 *mm/s* and decremented 1 *mm/s* per line with an actuation speed of 17.0 *μm/s.* The software that accompanies the microscope used to take these pictures allows for measurements in pixels and subsequent conversion to mm. Scale bar: 2mm.

**Supplementary Figure S5.**
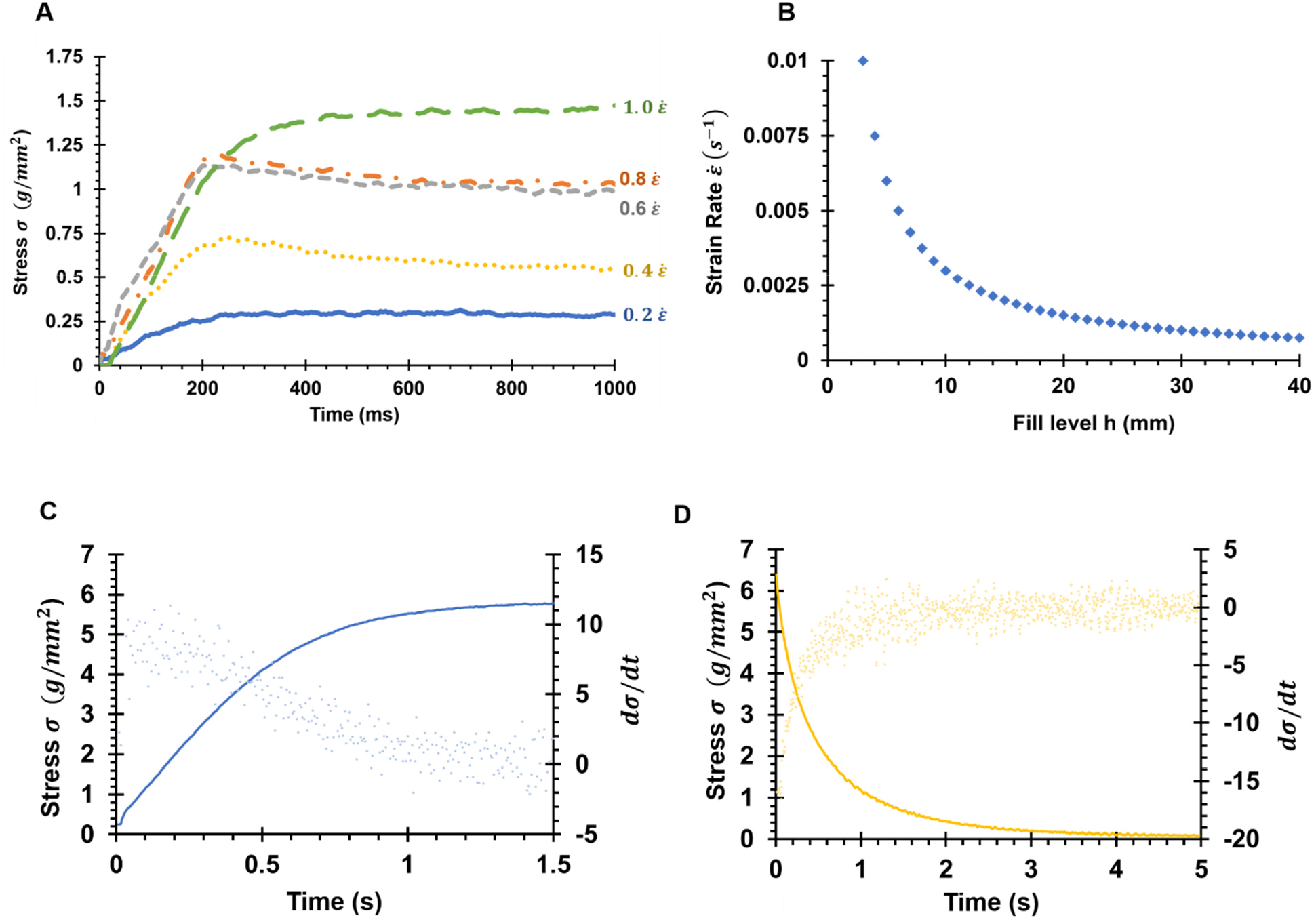
The viscoelasticity of NGM. The compressive strain rate is directly proportional to the equilibrium stress in the liquid NGM during extrusion. (**A**) Graph showing how stress (J changes over time, for 5 different strain rates *ε̇* (*s*^−1^). NGM extruded at a constant actuation speed will result in increasing strain rates (see also Supplementary Fig. S5). (**B**) Strain rate varies with the NGM fill level in the syringe and the actuation speed. Study done with 30 *μm/s* actuation speed. The syringe used in the Parnon is considered full at 50 linear millimeters of NGM. Eq. SF4 describes the inverse relationship of strain rate (*ε̇*) to fill level (*h*) given a constant actuation speed (v). *Eq. SF*4: 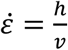. The values for strain rate (ε̇) are unique to the syringe used in the Parnon, but the trend is expected to apply in all cases of actuation pressure extrusion. (**C**): Comparison of increasing *σ* (continuous line, left y axis) and *dσ/dt* (scatter plot, right y axis) over time (stress build up). When *dσ/dt* reaches 0, the inflection point has been reached. The time it takes for the inflection point to be reached is *t_eq._*. (**D**): Comparison of decreasing (J (continuous line, left y axis) and *dσ/dt* (scatter plot, right y axis) over time (stress relaxation). When *dσ/dt* reaches 0, the inflection point has been reached. The time it takes for the inflection point to be reached is *t_r_*. It was found that *t_eq_*. = *t_r_* = ∼600 ms at *ε̇* = 1.1. Panels C and D: Compressive results using a 5mL, 12mm ID syringe and a 404 ID nozzle.

**Supplementary Figure S6:**
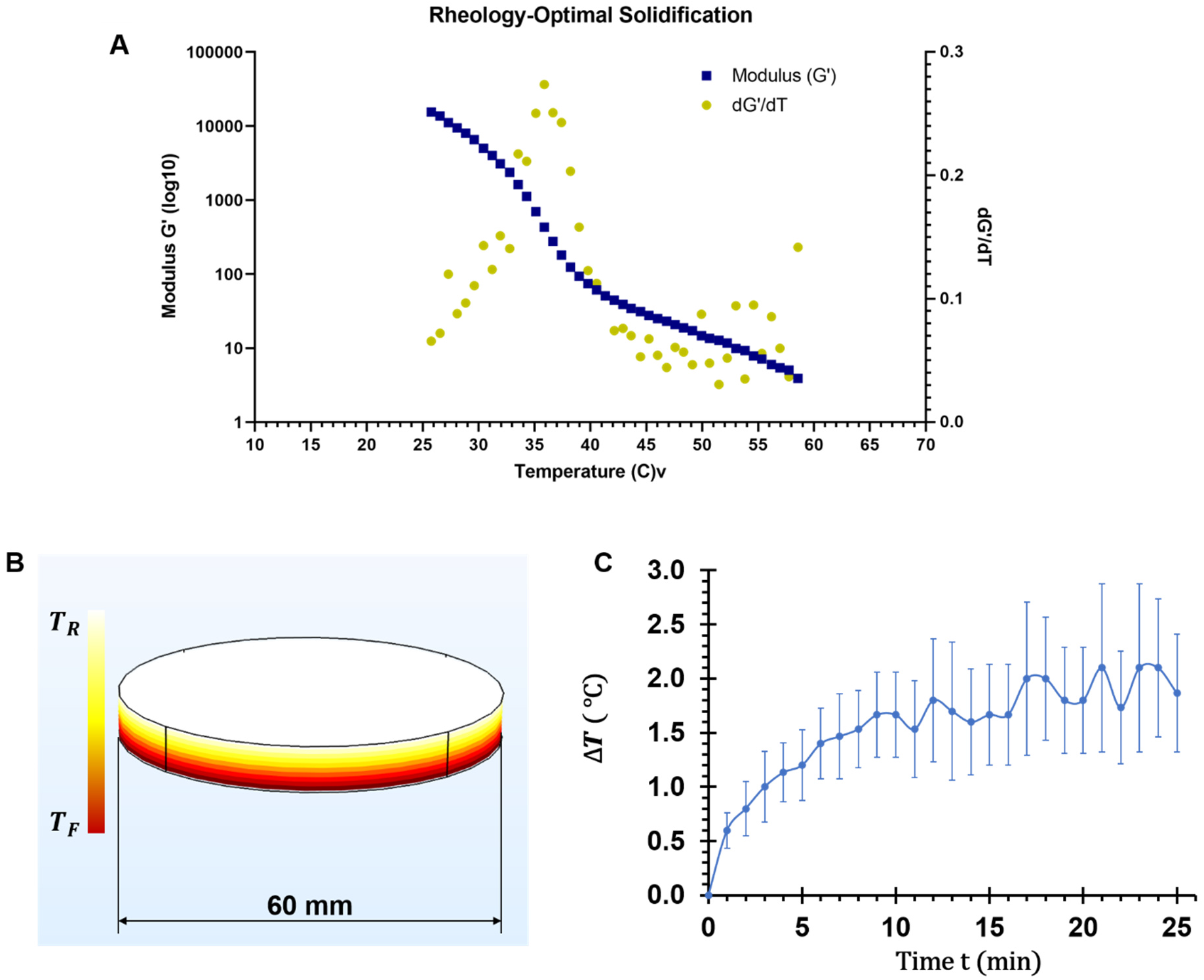
NGM solidification rheology, and heat flux model of the substrate. (**A**) Graph shows the storage modulus G’ (blue squares, left y axis) and rate of change dG’/dT (green circles, right y axis) as a function of temperature. The highest value of dG’/dT corresponds to 35.9 °C, meaning that at this temperature solidification occurs the fastest. Rheology experiment performed on a TA DHR2 Rheometer at a cooling rate of 5 °C /min and an angular frequency of 1 rad/s. (**B**) COMSOL visualization of the temperature gradient across the plotting medium (plotting medium cylinder volume: 20 mL). The cooling source is the heat flux (Q) provided by the Peltier device beneath the substrate. The temperature drop (Δ*T*) is defined as Δ*T* = *T_R_* - *T_F_*, where *T_R_* = 25 °C (estimated room temperature) and *T_F_* is dependent on the effective heat flux (Q). (**C**) Actual temperature drop over time, with mean of three independent experiments and standard error bars. The cooling levels out at a Δ*T* of ∼2.0 °C. Using Eq. F9 our Peltier device is achieving an effective heat flux of 182 *mW/cm*^2^ or ∼5.15 *W* over the entire 60 *mm* petri dish. Eq. F9 describes the linear relationship between Temperature Reduction (Δ*T*, in °C) and Heat Flux (Q, in (*mW/cm*^2^). *Eq. F9: ΔT=0.011Q*

**Supplementary Figure S7:**
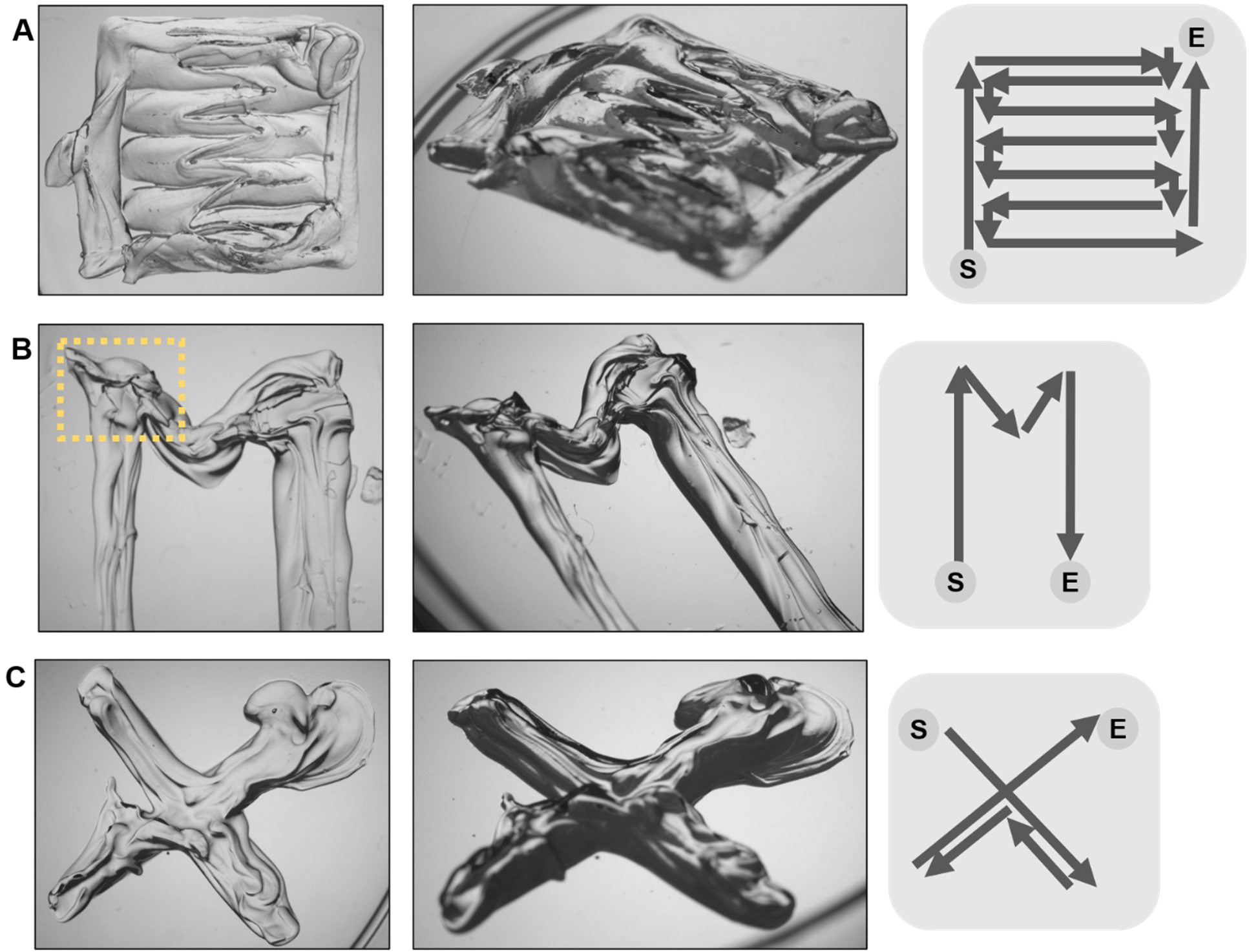
Parnon-printed structures and printing paths. These structures were printed with a larger diameter nozzle (400 *μm* - 1 *mm* ID) than the structures in Fig 5. (A) NGM pad, printed to test Parnon’s ability to create continuous sheets. (B) M-shaped print, to test Parnon’s ability to lay NGM lines in acute (<90°) angles, yellow frame indicates reduced printing quality area. (C) A cross-shaped print, to test Parnon’s ability to print cross-shaped designs, without having to pause extrusion, relocate, and restart extrusion. All panels: Left: top view; middle: perspective view; right: schematic of printing path; S: start, E: end.

**Supplementary Figure S8:**
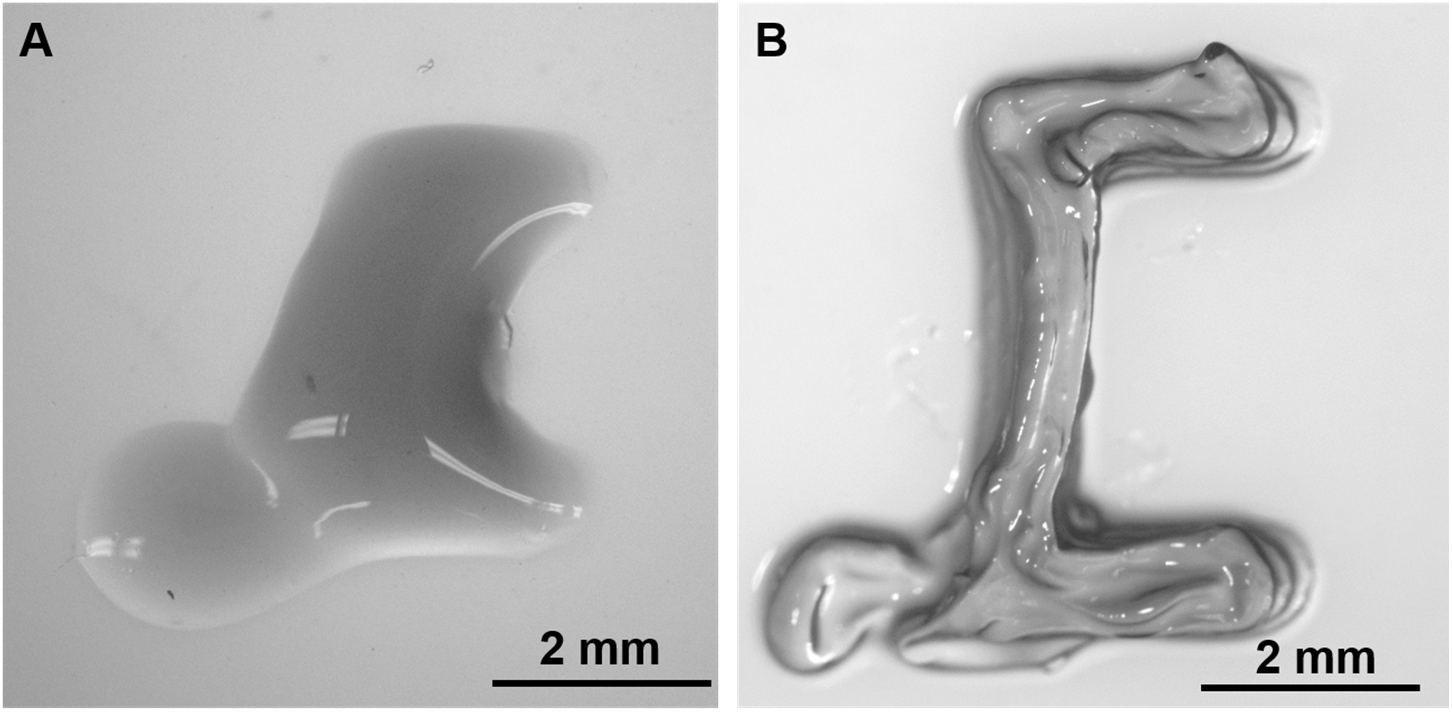
Printed part quality comparison with respect to plotting medium usage. (**A**) C-shaped part printed without plotting medium, exhibiting extensive liquid spreading, top view. (**B**) Same C-shaped design printed using plotting medium, with minimal liquid spreading, top view. This is the same C-shaped part as in Figure 5A & 5B.

**Supplementary Figure S9:**
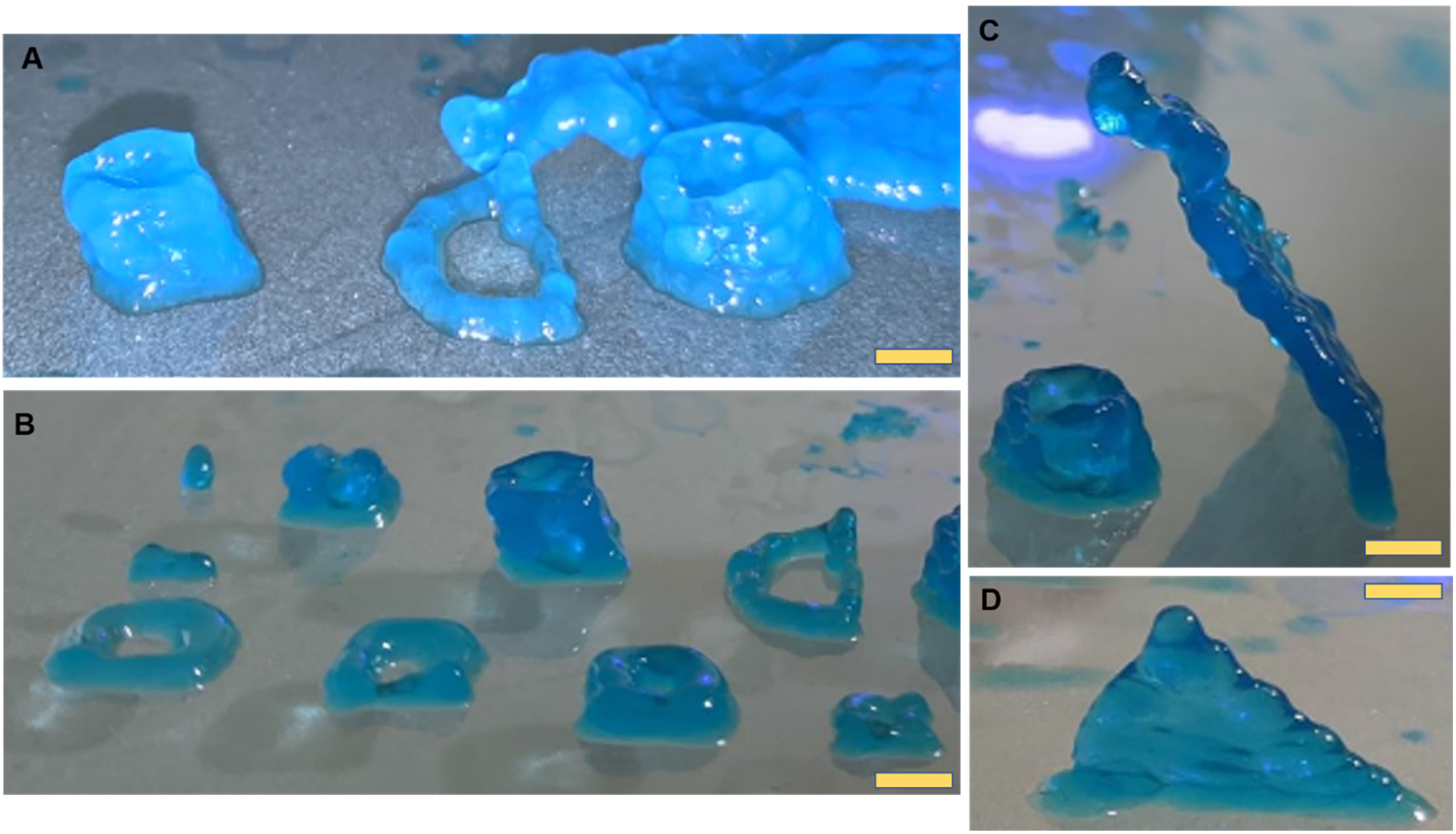
Printing attempts for multi-layer NGM parts. (**A**) Examples of 1- to 4-layer cylindrical structures, scale bar: 5mm. (**B**) Examples of 1- to 5-layer cylindrical structures, front row printed in the absence of plotting medium, scale bar: 6mm. (**C**) A 5-layer cylinder and a 14-layer triangular tilted wall (side view), scale bar: 6mm. (**D**) A12-layer triangular wall similar with the one in C, tilted forward (front view), scale bar: 3mm. Blue dye: food color (Americolor, USA).

## Supplementary Information

A full list of the major parts and items used for the customization of Parnon printer is provided in the Appendix. The Arduino code is available upon request.

### i. Preparation of 3D printed structures for C. elegans experiments

After printing, the 3D parts are removed from the plotting medium and are placed on an unseeded NGM plate. If they are to be used immediately, i.e. in the next 24 hours, they are rinsed five times with deionized water, then are left to dry off excess moisture, in sterile conditions. If they are to be used in longer than 24 hours, then they are washed with 5% sodium hypochlorite, for 15 min on an orbital shaker, followed by five rinses with deionized water, 10 min on the orbital shaker, each. This precaution is necessary since the parts are not printed in sterile environment. Next, the plates are left to dry off excess moisture in sterile conditions. The plates can be sealed with parafilm and stored in 4°C for several days, in which case they need to reach room temperature before use. If the plates are used after a few weeks, some evidence of crystallization might be noticeable in the plate NGM (see for example Fig. 6D and in less extent 6E), but the overall suitability is not compromised.

### ii. Actuation pressure

The linear actuator (Fig. 2 Part Ci) is a NEMA-8 captive with a 1: 1 step stroke of 3 *μm*. We use a 1: 4 step ratio for an effective step stroke (*l_st_*) of 0.75 *μm*. The actuator has a 38 *mm* total stroke length and a volume capacity of 2,470 *mm*^3^. A typical part design will utilize a volume of 50 *mm*^3^. Parnon’s capacity is roughly 50*x* the average print volume. It is powered by a 40V power supply through a TB6600 Stepper Motor Driver. The motor driver has microstepping capability, a 1: 2 step stroke is then 1.5 *μm*. The linear actuator is rated for to run at 0.49 *A*, the motor driver is set to 0.5 *A*. This linear actuator was chosen because it has the highest stroke precision the manufacturer could offer. An early version of Parnon utilized a NEMA-14 linear actuator that had a lower stroke precision. A higher stroke precision results in higher quality prints. The parts in Figs. 5C, 5D, 5F, and 6 are printed with the early version of Parnon, wearing NEMA-14; the parts in Figs. 5A, 5B, 5E, 7, and 8 are printed with the recent version of Parnon, featuring NEMA-8.

The Arduino card communicates with the motor driver by sending 5 *V* pulses in a square wave. A user defined delay time (*t_d_*) is how long the pulse is sustained for. Each pulse equates to one step taken by the linear actuator. A total print time (*t_pr_*) is derived during the gCode writing process. The total number of steps, or pulses, during a print control the extruded volume of NGM. A theoretical extruded volume (*V_ex_*.) can then be calculated.

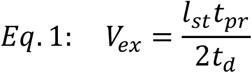

By varying *t_d_*, *V_ex_* can be controlled. A lower *t_d_* will result in a more fluent stepping process in the actuator and higher *V_pr_*.

The gCode defines the layer height (*h*_l_) and the print path length (*l_pr_*). Using the nozzle ID (*w_i_*), a theoretical print volume (*V_pr_*) can be calculated.

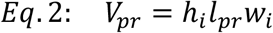

We refer to the designed ratio of the print as the ratio of *V_ex_* to *V_pr_*. Despite the Parnon’s several features meant to limit the amount of liquid spreading, some amount of liquid spreading still occurs. A print quality comparison between prints that resulted while the liquid spreading mitigations were active *vs* when the plotting medium and cooling effect were not used is presented in Supplementary Figure 4.

The amount of liquid spreading the fully functioning Parnon experiences was studied so the actual width of a line of printed NGM becomes predictable and designable. Results of the line width experiment are shown in Supplementary Figure 1. Four 20 *mm* long lines of NGM were printed, with 5 *mm* transitions. Two experiments, accelerating and decelerating, were repeated 3 times each. A relationship between the designed ratio (*V_ex_/V_pr_*) and the actual ratio (*w_l_/w_i_*) of the line width (*w_l_*) to the nozzle ID (*w_i_)*. A designed ratio of ∼0.9 equates to an actual ratio of 1.0 (Fig. 3).

We control the designed ratio by varying *t_d_* in *Eq.* 1, as the variables in *Eq.* 2 are set by hardware and the previously written gCode and thus more difficult to vary. By setting *V_ex_/V_pr_* = 0.9 we expect *w_l_/w_i_* = 1, or, the final line width after liquid spreading will be equal to the inner diameter of the nozzle.

### iii. NGM compressive viscoelastic response

NGM in the 9.11mm ID syringe used in the latest version of the Parnon was tested at strain rates starting at *ε̇* = 0.2, incrementing by 0.2 five times ending at *ε̇* = 1.0 (Fig. 4 Part A). *σ_eq_*. ranged from ∼0.25 *g/cm*^2^ at *ε̇* = 0.2 to ∼1.5 *g/cm*^2^ at *ε̇* = 1.0. Strain rates decrease inversely as the fill level in the syringe increases for a constant actuation speed [Suppl. Fig. 5]. In the 9.11mm ID syringe *t_eq._* = ∼300 *ms*.

NGM in the 12 mm ID syringe used in the early versions of the Parnon was tested 4 times. We found that *t_eq_*. ≅ 600ms at a *ε̇* = 1.1. (Fig. 4 Part Bi). In the 12mm ID syringe *t_eq_*. = ∼600ms. We believe the larger ID of this syringe caused the higher *t_eq_*. and *σ_q_*. when compared to the lower syringe ID (9.11mm) in the current version of the Parnon.

It was found that *t_eq_*. = *t_r_*. The stress in the NGM during extrusion will increase with both the syringe ID and the strain rate (*ε̇*). The time it takes to reach the inflection point increases with the syringe ID but is effectively constant across multiple strain rates. In practice, we found that *t_eq_*.= 700ms with the 9.11mm ID syringe results in the most accurate extrusion timing.

The time *t_eq_* it takes for the compressive stress *σ*_eq_ in the NGM to reach a high enough level to begin pushing a relatively viscous liquid through a nozzle with a small ID (254 *μm* - 400 *μm*) increases with strain rate. As part of the work done on optimizing Parnon there is a feature written in the code that automatically uses *t_eq_* to build up to *σ_eq_* immediately before beginning extrusion during a print. With this addition the Parnon begins extruding NGM at the right time. Before, the NGM often extruded late as time had to pass for *σ_eq_* to be reached. We speculate that the NGM near the nozzle where the resulting curves rose above *σ_eq_* before approaching it (*ε̇* = 0.4, 0.6. 0.8) were more viscous and therefore required more stress to be extruded. In addition, *σ_eq_* for *ε̇* = 0.6, 0.8 are relatively close in value. We believe this occurs because of differing viscosity conditions in the NGM during extrusion.

### iv. gCode and software communication

Modifying the hardware of an existing 3D printer compromised the communication the printer had with the original print head. To resolve this issue, we introduced a limit switch, an Arduino card, and a stepper motor driver.

Before every print, a few lines of gCode instruct the Parnon print head to hit the limit switch, thus signifying to the software that the print has begun.

An Arduino Uno board connected to a Stepper Motor Driver which in turn is connected to the linear actuator. A custom Microsoft Excel program that processes raw cartesian coordinates and head speed values outputs three essential printer inputs:

a. A gCode file that the DOBOT MOOZ-2 will interpret into XYZ axis actuation commands. The gCode file is loaded on a USB external drive or a micro SD card and placed in the appropriate slot on the Parnon.
b. The time between the custom print head hitting the limit switch and beginning actuation pressure. We refer to this value as delay one (*delay*1) and it is measured in milliseconds.
c. An array of step delays (measured in milliseconds), steps to take and the relative volume (Designed Ratio Fig. 3 Part B). The step delay (*t_d_*) and number of steps (*n*) which correspond to a designed ratio of ∼0.90 are inputted through the Arduino serial port.

Outlining the part is currently done in a custom Microsoft Excel program. Each coordinate XYZ that the print head will travel to during the print is individually assigned to a row in order. Upon executing the custom Microsoft Visual Basic program, the coordinates are processes and written into a gCode file. The gCode is processed by the existing firmware from DOBOT.

Time needs to pass between the print head hitting the limit switch and the point at which extrusion should begin. This time is referred to as delay one (*delay*1), and depending on the part design varies around 2.4 *s*. The viscoelastic response NGM has to compressive stress requires a significant amount of time (*t_eq_*. = 700 *ms*) to pass before sufficient stress has been reached for extrusion to commence. *t_eq_*. is included in *delay*1 and is referred to as *strain* in the Arduino Code. Section iv of Supplementary Information refers in detail to NGM’s response to compressive stress.

The total print time (*t_pr_*) is derived in the Excel program as a result of dividing the path length (*l_pr_*) by the head speed (*v_h_*). *l_pr_* and *v_h_* are defined in the gCode. Depending on the delay time between each step (*t_d_*), the number of steps for the actuator to take (*n*) so it stops extruding when the print stops can be calculated.

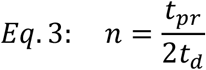

Depending on the distance each actuator step is (*l_st_*), discussed in more detail in the next section, there will be a specific calculatable volume for each selection of *t_d_*. Multiplying the cross-sectional are of the syringe (A_s_ = 65 *mm*^2^) by the distance the actuator will travel during the print allows the theoretical total extruded volume (*V_ex_*) to be calculated.

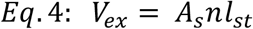

Comparing this theoretical volume to the total print volume (*V_pr_*) gives us what we refer to as the designed ratio (*V_ex_/V_pr_*). How this value compares to what can actually be observed and is discussed in detail in the next section.

### v. Plotting medium

A COMSOL Multiphysics simulation was ran on 20 *mL* of plotting medium in a 60 *mm* diameter cylinder (Fig. 5 Part C). A negative heat flux was placed on the bottom of the geometry to model the cooling effect of the Peltier device. The relationship between the heat flux (*Q*, *mW*/*cm*^2^) and the temperature drop across the plotting medium (Δ*T*, °C) is described by *Eq.* 5.

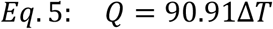

It was determined experimentally that after ∼20min, Δ*T* = ∼2.0 °C (Fig. 5 Part D). As a result of the COMSOL modeled relationship shown in Eq. 5, *Q* = 182 *mW/cm*^2^. Over the whole area of the plotting medium (2.83 *cm*^2^) the energy exiting the plotting medium is 5.15 *W*. The Peltier device is rated at 6 *W*. The Parnon’s cooling system operates at 86% efficiency. During a print, the NGM is being extruded into plotting medium that is ∼13 °C less than the fastest solidification temperature. A dynamic heat transfer model of NGM cooling in the plotting medium was not developed. NGM is extruded at ∼65 °C, however, ∼29 °C above the fastest solidification temperature. Assuming that the thermal conductivities of NGM and the plotting medium are equal, then NGM would solidify the fastest if the plotting medium was also ∼29 °C less than the fastest solidification temperature. There is another factor to consider, however, FDM relies on the previous layer remaining liquidous until the next layer is able to fuse. If the NGM solidifies too quickly the layers will not fuse together resulting in failed prints. While not quantified, we speculate that the limited ability of the Parnon’s substrate cooling system is beneficial to the overall printing capability.

## Appendix

*List of Parnon major parts and items used for customization*

- DOBOT MOOZ-2 machine, Shenzhen, China
- 12V DC Power Supply, 5V Power Supply
- Arduino SD Cardboard, Arduino UNO
- Heating Cartridge: Insertion Heater with Internal Temperature Sensor, for 3/8” Hole, 120V AC, 3” Long Heating Element, 200W, McMaster-Carr, USA
- Heating Controller: Programmable Temperature Controller, for Type K Thermocouple, McMaster-Carr, USA
- Limit Switch: Monoprice Maker Select 3D Printer 13860 Endstop Limit Switch, Amazon, USA
- Linear Actuator: NEMA-8 captive, HeydonKerk, USA
- Microstep Driver: TB6600 4A 9-42V Stepper Motor Driver CNC Controller, Amazon, USA
- Motor Driver: EasyDriver V4.5, SparkFun, USA
- Peltier Devices: Northbear Thermoelectric Cooler Peltier Refrigeration Cooling TEC System Kits Double Fan DIY +Power Supply, NorthBear, USA
- Thermal Compound: ARCTIC MX-4 - Thermal Paste for Coolers, Heat Sink Paste

### Details on the Parnon customized print head

The customized print head includes the following parts (Fig. 4C): **i)** Bipolar stepper motor of the linear actuator, to provide the actuating force with a 40V input, controlled with Arduino. **ii)** 3D-printed custom connector (FormLabs Form2 3D printer, clear resin v4), used to house the stroke arm (iii) and to connect the motor (i) to the heat sink (v). **iii)** Stroke arm of the linear actuator with a 1.5” total actuation range and a 1:1 stroke of 3 μm. **iv)** Heating element, 3” length, 3/8” diameter, 200W, set to 65 °C, used to provide the temperature required to keep the NGM in a liquid state. **v)** Custom heat sink designed to transfer heat from the heating element (iv) to the syringe (viii). **vi)** Connector (FormLabs Form2 3D printer, tough resin v5), used to connect the stroke arm (iii) to the plunger (vii). **vii)** Glass syringe plunger which has the custom connector (vi) adhered on the inside. **viii)** Glass syringe that houses the liquid NGM. **ix)** Metal luer-lock nozzle. **x)** Copper wire heat induction system the prevents NGM solidification in the nozzle (ix).

## Notes

### Competing Interest Statement

The authors have declared no competing interest.

